# Brain perivascular space imaging across the human lifespan

**DOI:** 10.1101/2022.01.25.475887

**Authors:** Kirsten M. Lynch, Farshid Sepehrband, Arthur W. Toga, Jeiran Choupan

## Abstract

Enlarged perivascular spaces (PVS) are considered a biomarker for vascular pathology and are observed in normal aging and neurological conditions; however, research on the role of PVS in health and disease are hindered by the lack of knowledge regarding the normative time course of PVS alterations with age. To this end, we characterized the influence of age, sex and cognitive performance on PVS anatomical characteristics in a large cross-sectional cohort (∼1400) of healthy subjects between 8 and 90 years of age using multimodal structural MRI data. Our results show age is associated with wider and more numerous MRI-visible PVS over the course of the lifetime with spatially-varying patterns of PVS enlargement trajectories. In particular, regions with low PVS volume fraction in childhood are associated with rapid age-related PVS enlargement (e.g., temporal regions), while regions with high PVS volume fraction in childhood are associated with minimal age-related PVS alterations (e.g., limbic regions). PVS burden was significantly elevated in males compared to females with differing morphological time courses with age. Together, these findings contribute to our understanding of perivascular physiology across the healthy lifespan and provide a normative reference for the spatial distribution of PVS enlargement patterns to which pathological alterations can be compared.

## Introduction

The brain waste clearance system consists of a network of vasculature that plays a critical role in the removal of toxic substrates from the brain to maintain tissue homeostasis. A major component of the waste clearance system includes perivascular spaces (PVS), which consist of tubular, interstitial fluid-filled cavities that surround small penetrating vessels in the brain parenchyma (Wardlaw et al., 2020). PVS provide a low-resistance pathway to facilitate fluid exchange (Bedussi et al., 2018), where they accommodate the influx of cerebrospinal fluid (CSF) and energy substrates in the peri-arterial space and the drainage of interstitial fluid and metabolic waste through the peri-venous space (Bacyinski et al., 2017; Iliff et al., 2013, 2012). It was recently demonstrated in animal models that pathological PVS enlargement is associated with reduced waste clearance functionality (Xue et al., 2020) and can thus impede the removal of toxic metabolites, including amyloid beta (Aβ), and render the brain susceptible to neurological damage (Keable et al., 2016). Because several neurological conditions are characterized by dysfunctional waste clearance (Sweeney et al., 2018; Troili et al., 2020; Wardlaw et al., 2020), an understanding of the factors that contribute to PVS alterations can provide insight into disease pathogenesis.

Over the past decade, technological advancements in MRI acquisition and data processing have facilitated the study of PVS characteristics in health and neurological disease. Recent evidence from MRI studies in humans have shown significantly increased PVS visibility in the brains of patients with neurological conditions, including cerebrovascular disease (Charidimou et al., 2017, 2013; Doubal et al., 2010; Martinez-Ramirez et al., 2013; Potter et al., 2015), traumatic brain injury (Opel et al., 2018), stroke (Charidimou et al., 2013), Alzheimer’s Disease (AD) (Banerjee et al., 2017b), mild cognitive impairment (Sepehrband et al., 2021) and Parkinson’s disease (Donahue et al., 2021). While typically considered a biomarker of vascular neuropathology (Doubal et al., 2010; Mestre et al., 2017; Potter et al., 2015), enlarged PVS are also observed in healthy, cognitively normal individuals. Increased PVS visibility is a prominent feature of advancing age (A Laveskog et al., 2018; Zhu et al., 2011) and recent evidence has shown that enlarged PVS are also observed in typically developing adolescents (Piantino et al., 2020) and young adults (Barisano et al., 2020), albeit at much lower levels. However, the precise trajectory of PVS alterations as a consequence of age across the normative lifespan has yet to be fully described. Because the highest risk factor for the development of neurodegenerative disease is advancing age (Hou et al., 2019), age-related alterations to the components that facilitate the elimination of waste may render the brain more vulnerable to neurodegenerative pathology. Therefore, characterization of the time course of PVS alterations in the normal human brain can provide insight into the evolution of waste clearance mechanisms and can offer a benchmark from which pathological PVS enlargement can be differentiated from that attributed to normal, healthy aging.

PVS have been observed in several regions of the brain, including the centrum semiovale of the white matter, basal ganglia (BG), hippocampus and midbrain structures (Potter et al., 2015). In particular, PVS enlargement within the white matter and BG are differentially associated with neuropathology. Vasculopathies, such as cerebral small vessel disease (CSVD) and hypertension, are preferentially associated with increased PVS visibility in the BG (Charidimou et al., 2017, 2013; Doubal et al., 2010; Martinez-Ramirez et al., 2013; Potter et al., 2015), while white matter PVS alterations are predominantly observed in patients with amyloidopathies (Banerjee et al., 2017a; Charidimou et al., 2015; Martinez-Ramirez et al., 2013; Ramirez et al., 2015), such as AD and cerebral amyloid angiopathy (CAA). The apparent spatial dependence of neuropathological PVS enlargement suggest the mechanisms that contribute to increased PVS visibility differ between the white matter and BG. The heterogeneous characteristics of MRI-visible PVS therefore suggest that the mechanisms that govern PVS dilations in health and disease may differ according to spatial location. A comprehensive understanding of regional patterns of PVS enlargement across the lifespan in the BG and subcortical white matter would provide granular insight into brain regions that may be particularly susceptible to age-related changes in waste clearance processes.

Much of the evidence of age-related changes to PVS burden come from studies that employ a visual rating scale to score the severity of PVS burden on select axial slices (Francis et al., 2019; Gutierrez et al., 2013; A. Laveskog et al., 2018; Yakushiji et al., 2014; Zhu et al., 2011, 2010). While this approach can be easily implemented in clinical settings, it does not objectively quantify PVS content and may fail to capture regional PVS heterogeneity across the brain. Recently, efforts that utilize automated methods have enabled quantification of global PVS features; however, studies of normative aging have largely focused on PVS total volume or the fraction of tissue volume occupied by PVS (volume fraction, VF) (Barisano et al., 2020; Huang et al., 2021). These approaches provide limited information regarding the anatomical characteristics of PVS and can be confounded by total brain volume and age-related atrophy, as previous studies have shown PVS volume is significantly correlated with intracranial volume (ICV) (Barisano et al., 2020; Huang et al., 2021). Therefore, utilization of multiple morphological and structural characteristics, such as cross-sectional diameter, solidity and count, can provide greater insight into the mechanisms that contribute to age-related alterations in PVS volume.

The goal of this study is to characterize the influence of age on regional PVS burden in the white matter and BG across the lifespan in a large cross-sectional cohort (∼1400) of typically developing and cognitively normal children, adults and the elderly between 8 and 90 years of age. Here, we visualize PVS using an automated processing workflow to identify and quantify PVS morphometric features from vesselness maps derived from multi-modal structural neuroimaging contrasts designed to enhance PVS visibility (Sepehrband et al., 2019). Features of PVS morphology, including PVS VF, count, mean cross-sectional diameter and mean solidity, were extracted and used to quantify the magnitude and timing of PVS alterations across the lifespan in the BG and white matter regions using growth models. Additionally, we explored the influence of sex, excess body mass and blood pressure on age-related alterations to PVS morphology. The results from this study will provide a normative reference for the spatial distribution and time course of PVS alterations across the lifespan, to which pathological deviations from the expected trajectory can be compared.

## Results

Age distributions for the HCP cohorts stratified by sex are provided in **Table 1**. Overall, age was not significant different between sexes (*t*(1387)=.79, *p*=.43); however females were significantly older than males in the HCP-YA cohort (*t*(403)=-4.07, *p*<.001). PVS were present throughout the white matter and BG in all 3 cohorts and showed increasing presence with age (**Fig. 1)**.

**Fig. 1.**
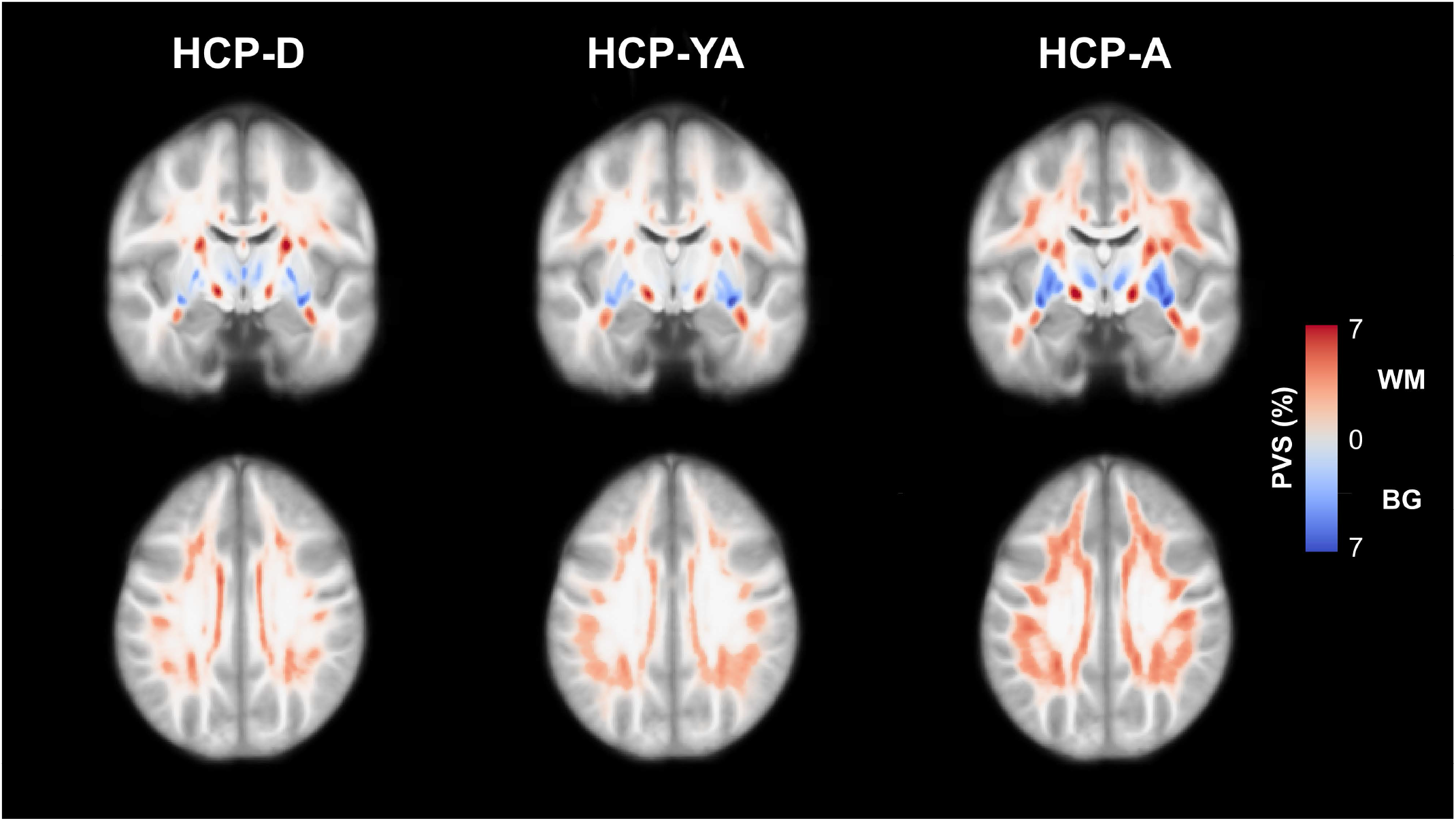
Maps of the PVS distribution in the HCP-D (n=471), HCP-YA (n=405) and HCP-A (n=516) cohorts. Normative PVS maps were generated by registering each subject map to a population-averaged template and then averaging PVS segmentations across each group. The results are presented as the percentage of overlapping PVS across each group within the subcortical white matter (red) and basal ganglia (blue). PVS is prevalent in all HCP cohorts, including children and adolescents, and the apparent PVS burden increases with age in both the white matter and basal ganglia.

**Table 1.** Age distribution of HCP cohorts stratified by sex.

| Dataset | *n* | Age |  | *t*-statistic |
| --- | --- | --- | --- | --- |
|  |  | Mean (SD) | Range |  |
| HCP-D | 471 | 14.36 (3.77) | 8.08-21.92 | -- |
| *Males* | 211 | 14.44 (3.57) | 8.08-21.83 | 0.43 |
| *Females* | 260 | 14.29 (3.93) | 8.17-21.92 |  |
| HCP-YA | 405 | 28.73 (3.78) | 22-36 | -- |
| *Males* | 179 | 27.89 (3.64) | 22-36 | -4.07*** |
| *Females* | 226 | 29.40 (3.78) | 22-36 |  |
| HCP-A | 513 | 56.35 (13.65) | 36-89.75 | -- |
| *Males* | 215 | 56.69 (13.74) | 36.00-89.75 | 0.47 |
| *Females* | 298 | 56.11 (13.59) | 36.17-87.75 |  |
| Combined | 1389 | 34.17 (20.07) | 8.08-89.75 | -- |
| *Males* | 605 | 33.70 (20.16) | 8.08-89.75 | 0.78 |
| *Females* | 784 | 34.54 (20.00) | 8.08-87.75 |  |

SD = standard deviation; \*\*\* $p < .001$ \*\* $p < .01$ \* $p < .05$

### Age-related PVS alterations in the basal ganglia

Cohort-stratified multiple regression analyses show PVS morphological features are significantly associated with age after controlling for covariates in the HCP-A group only (**Table 2**). Age is positively associated with PVS volume fraction (VF) (*p*<.0001), PVS count (*p*<.0001), and mean PVS cross-sectional diameter (p<.0001) and negatively associated with mean PVS solidity (p<.0001) in the HCP-A cohort **(*SI Appendix*, Fig. S1**). In the combined lifespan analyses, a quadratic regression best explained the age-related variance in PVS morphology (**Table 3**). The relationship between age and PVS VF, mean PVS diameter and PVS count were described with a convex curve (**Fig. 2A-C**) and the relationship between age and mean PVS solidity were described with a concave curve (**Fig. 2D**). The minimum PVS volume fraction and PVS count are estimated at 14±3 years and 43±2 years of age, respectively, and increase thereafter, while the maximum PVS mean solidity is estimated at age 10±9 years and decreases nonlinearly. Across all features, the most rapid feature differences with age occur in the aging cohort (**Fig. 2E**).

**Table 2.** Association between PVS morphological features and age stratified by cohort

| Variable | Cohort | White matter |  |  | Basal ganglia |  |  |
| --- | --- | --- | --- | --- | --- | --- | --- |
|  |  | *B* | Std Error | *t* | *B* | Std Error | *t* |
| VF | HCP-D | $4.32 \times 10^{-4}$ | $3.65 \times 10^{-5}$ | 3.93*** | $2.11 \times 10^{-5}$ | $5.08 \times 10^{-5}$ | 0.41 |
| | HCP-YA | $1.82 \times 10^{-4}$ | $4.85 \times 10^{-5}$ | 3.75** | $4.45 \times 10^{-6}$ | $7.22 \times 10^{-5}$ | 0.06 |
| | HCP-A | $1.78 \times 10^{-4}$ | $1.26 \times 10^{-5}$ | 14.07*** | $3.63 \times 10^{-4}$ | $1.99 \times 10^{-5}$ | 18.29*** |
| Count | HCP-D | 32.58 | 4.59 | 7.09*** | -1.37 | 0.75 | -1.81 |
|  | HCP-YA | 15.19 | 5.34 | 2.85** | 0.67 | 0.73 | 0.91 |
|  | HCP-A | 1.18 | 1.41 | 0.84 | 0.95 | 0.16 | 5.78*** |
| Diameter | HCP-D | $6.69 \times 10^{-4}$ | $3.97 \times 10^{-3}$ | 0.17 | $3.36 \times 10^{-3}$ | $1.95 \times 10^{-3}$ | 1.72 |
| | HCP-YA | $1.52 \times 10^{-2}$ | $5.41 \times 10^{-3}$ | 2.81* | $6.18 \times 10^{-3}$ | $3.22 \times 10^{-3}$ | 1.92 |
| | HCP-A | $1.18 \times 10^{-2}$ | $1.50 \times 10^{-3}$ | 7.88*** | $1.06 \times 10^{-2}$ | $8.81 \times 10^{-4}$ | 12.08*** |
| Solidity | HCP-D | $-3.76 \times 10^{-4}$ | $3.33 \times 10^{-4}$ | -1.13 | $1.42 \times 10^{-4}$ | $3.40 \times 10^{-4}$ | 0.42 |
| | HCP-YA | $-1.18 \times 10^{-3}$ | $3.66 \times 10^{-4}$ | -3.22** | $9.59 \times 10^{-5}$ | $4.15 \times 10^{-4}$ | 0.23 |
| | HCP-A | $-3.68 \times 10^{-4}$ | $9.02 \times 10^{-5}$ | -4.08*** | $-1.02 \times 10^{-3}$ | $9.55 \times 10^{-5}$ | -10.70*** |

The partial correlation coefficients for the influence of age on PVS morphological features after controlling for sex and scanner type for the white matter and BG stratified by cohort. Age-related changes to white matter PVS morphology were observed in all cohorts, while BG PVS morphological changes with age were only observed in the aging cohort. \*\*\* $p < .001$ \*\* $p < .01$ \* $p < .05$

**Table 3.** Quadratic model coefficients that relate PVS morphological features with age across the lifespan in white matter and basal ganglia

| Coefficient |  | White matter |  |  |  | Basal ganglia |  |  |  |
| --- | --- | --- | --- | --- | --- | --- | --- | --- | --- |
|  |  | *B* | Std Error | *t* | *R*<sup>2</sup> | *B* | Std Error | *t* | *R*<sup>2</sup> |
| VF | Age | 1.96 x 10<sup>-4</sup> | 4.78 x 10<sup>-6</sup> | 40.93*** | 0.55 | -1.08 x 10<sup>-4</sup> | 3.40 x 10<sup>-5</sup> | -3.18** | 0.37 |
|  | Age<sup>2</sup> | -- | -- | -- |  | 3.90 x 10<sup>-6</sup> | 3.97 x 10<sup>-7</sup> | 9.83*** |  |
| Count | Age | 39.26 | 2.45 | 16.03*** | 0.30 | -2.37 | 0.36 | -6.55*** | 0.03 |
|  | Age<sup>2</sup> | -0.32 | 0.03 | -11.27*** |  | 2.94 x 10<sup>-2</sup> | 4.23 x 10<sup>-3</sup> | 6.95*** |  |
| Diameter | Age | 1.09 x 10<sup>-2</sup> | 5.40 x 10<sup>-4</sup> | 20.18*** | 0.23 | 2.04 x 10<sup>-3</sup> | 1.35 x 10<sup>-3</sup> | 1.51 | 0.29 |
|  | Age<sup>2</sup> | -- | -- | -- |  | 6.46 x 10<sup>-5</sup> | 1.58 x 10<sup>-5</sup> | 4.09*** |  |
| Solidity | Age | -1.27 x 10<sup>-3</sup> | 1.59 x 10<sup>-4</sup> | -7.96*** | 0.20 | 1.87 x 10<sup>-4</sup> | 1.79 x 10<sup>-4</sup> | 1.05 | 0.14 |
|  | Age<sup>2</sup> | 7.05 x 10<sup>-6</sup> | 1.86 x 10<sup>-6</sup> | 3.79*** |  | -9.56 x 10<sup>-6</sup> | 2.09 x 10<sup>-6</sup> | -4.57*** |  |

Quadratic regression fits for the PVS morphological features in subcortical white matter and basal ganglia. The quadratic model did not provide the best fit for the white matter PVS VF and mean diameter and age-related variance was better explained by a linear regression. \*\*\* p<.001 \*\* p<.01
\* p<.05

**Fig. 2.**
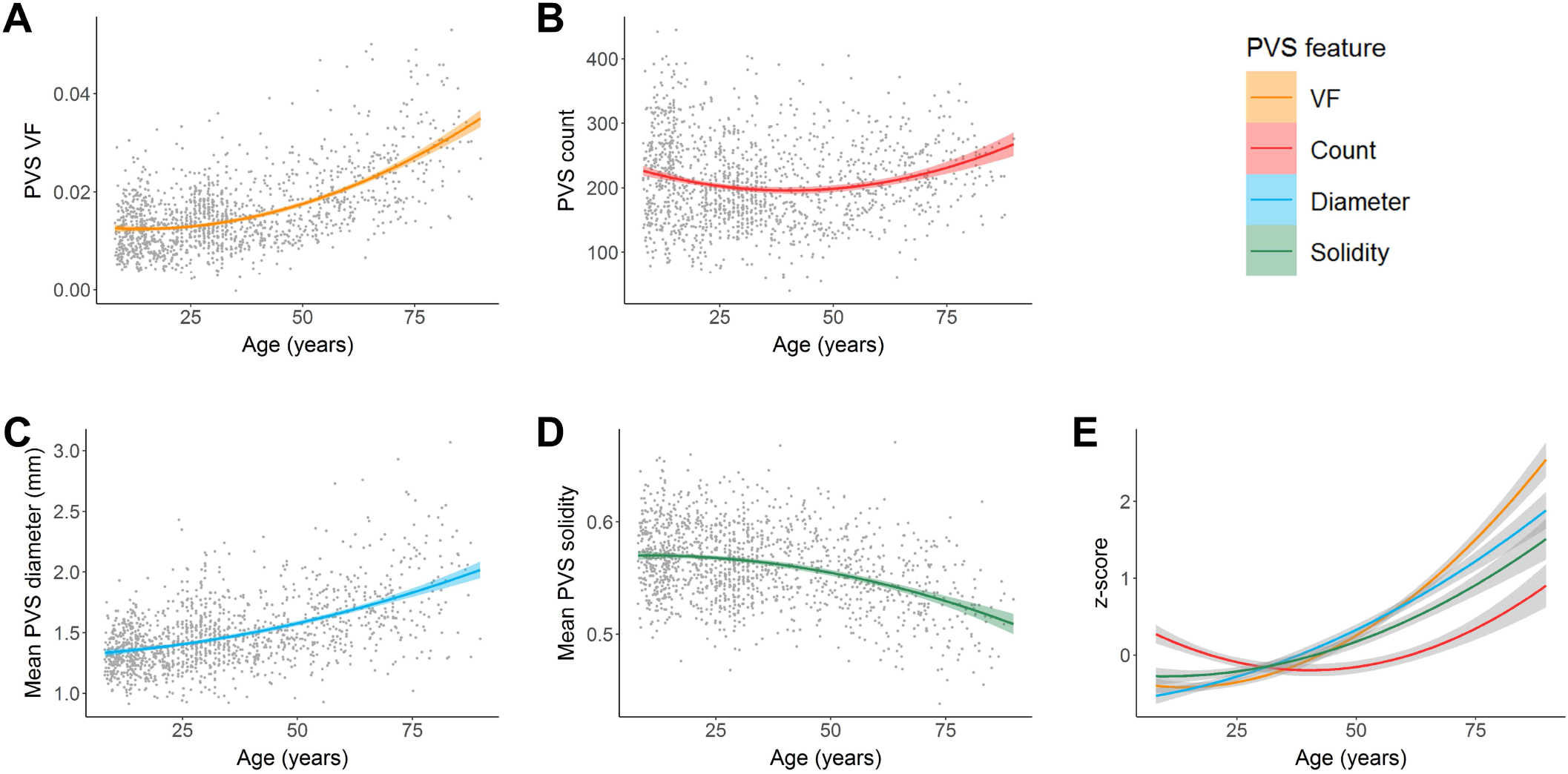
Associations between PVS morphology and age in the basal ganglia. A quadratic curve provided the best fit model to describe the relationship between age and (A) PVS volume fraction, (B) count, (C) mean diameter and (D) mean solidity. (E) PVS features in the BG were standardized (z-scores) and plotted on the same normalized y-axis to enable comparison of feature differences with age over the lifespan (x-axis). The mean PVS solidity curve was also inverted to show similarities and differences more easily. Within the BG, PVS volume fraction and mean PVS diameter undergo similar age-related trajectories, while mean PVS solidity and PVS count grow more slowly.

### Age-related PVS alterations in the total subcortical white matter

Cohort-stratified multiple regression analyses of PVS morphology in white matter show significant associations with age in multiple HCP cohorts (**Table 2**). PVS VF was significantly and positively associated with age in HCP-D (*p*<.0001), HCP-YA (*p*=.0002) and HCP-A (*p*<.0001). Age was significantly associated with increasing PVS count in HCP-D (*p*<.0001), but not HCP-A (*p*=.40). The relationship between age and PVS count in the HCP-YA (*p*=.0047) was reduced to non-significance following multiple comparison correction with a Bonferroni corrected threshold of p<.0042. Age was significantly associated with increased mean PVS diameter in HCP-A (*p*<.0001), but not HCP-D (*p*=.87) or HCP-YA following multiple comparison correction (*p*=.0053). Mean PVS solidity was significantly and negatively associated with age in HCP-A (*p*<.0001) and HCP-YA (*p*=.0014), but not HCP-D (*p*=.26). When combined across cohorts (**Table 3**), a linear regression account for the most age-related variance in PVS VF (**Fig. 3A**) and mean PVS diameter (**Fig. 3C**). The relationship between age and PVS count were best described by a concave quadratic model with the most rapid changes observed early in life and peaking at 61±2 years of age (**Fig. 3B**). Age-related reductions to PVS mean solidity were best described by a convex quadratic model that reaches stasis later in life with the estimated minimum solidity beyond the age range sampled (90±25 years) (**Fig. 3D**). Standardization of PVS features show all morphological characteristics contribute to the age-related increase in PVS VF with varying degrees (**Fig. 3E**).

**Fig. 3.**
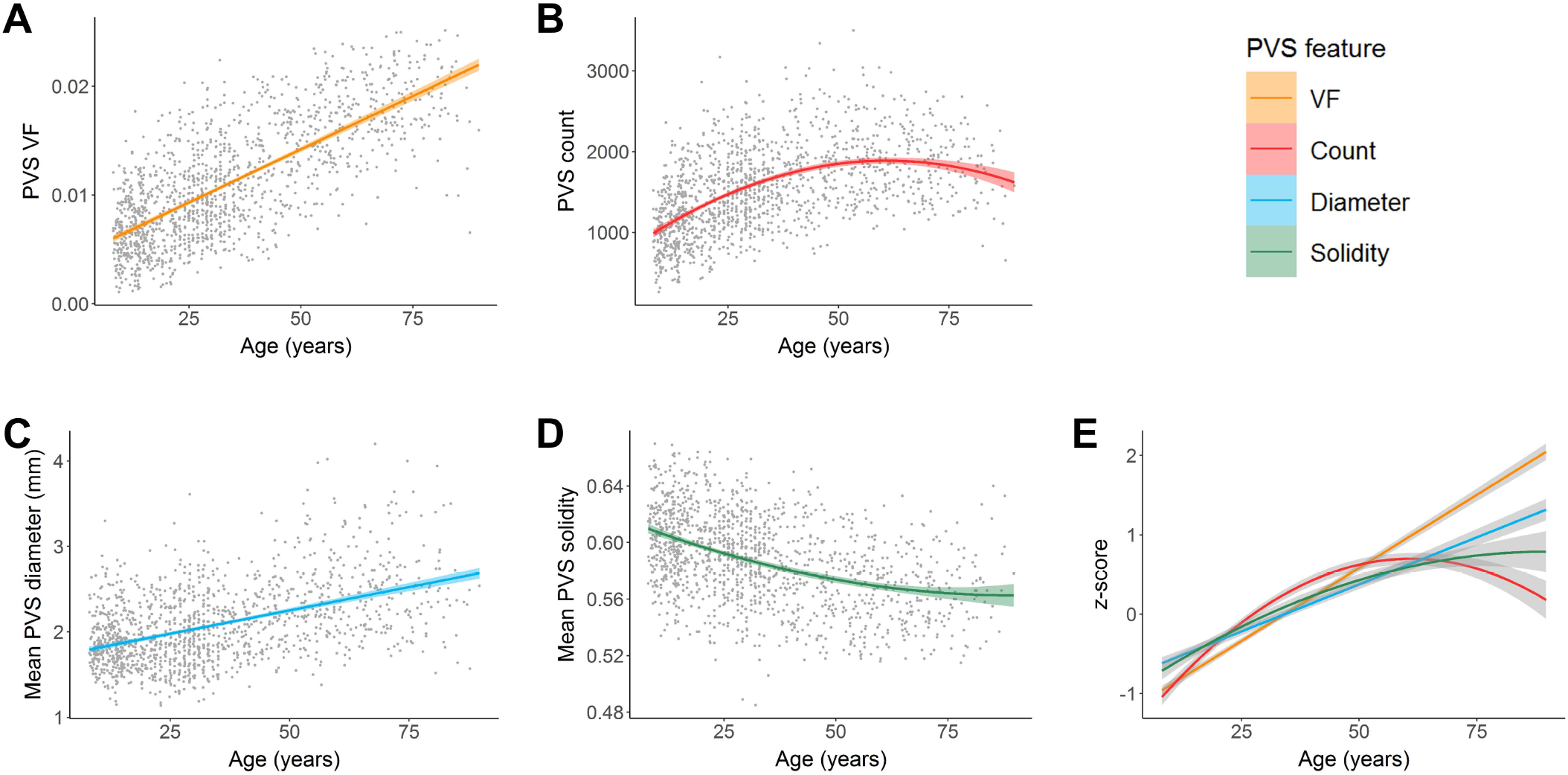
Associations between PVS morphology and age in the subcortical white matter. Age was positively associated with (A) PVS volume fraction, (B) count and (C) mean diameter and negatively associated with (D) mean solidity. Age-related changes to the PVS volume fraction and mean diameter were best explained with a linear model, while age-related changes to the PVS count and mean solidity were best represented with a quadratic model. (E) PVS features in the subcortical white matter were standardized (z-scores) and plotted on the same normalized y-axis to enable comparison of feature differences with age over the lifespan (x-axis). The mean PVS solidity curve was also inverted to show similarities and differences more easily. Within the white matter, PVS count, mean PVS diameter and mean PVS solidity undergo slower age-related changes compared to PVS VF.

### Age-related PVS alterations within regions of the subcortical white matter

The distribution of PVS burden across regions per HCP cohort is shown in **Fig. 4**. Within HCP-D, temporal white matter has the lowest PVS VF (*M*±*SD* = .0048±.0027), particularly within parahippocampal and entorhinal regions, followed by occipital (.0051±.0022), parietal (.0077±.0035) and frontal (.0071±.0044) structures. The limbic white matter has the highest PVS VF in HCP-D (.0143±.0047), specifically within bilateral isthmus and rostral anterior cingulate regions. Within HCP-YA, occipital white matter has the lowest PVS VF (.0062±.0028), following by temporal (.0076±.0034), parietal (.0100±.0042), frontal (.0119±.0047) and limbic (.0164±.0046) structures. This trend was also observed in the HCP-A cohort (occipital: .0099±.0039; temporal: .0132±.0045; parietal: .0159±.0051; frontal: .0186±.0052; limbic: .0217±.0053), with the lowest PVS VF in bilateral cuneus, parahippocampal and entorhinal regions and highest PVS VF in bilateral insula, rostral anterior cingulate and lateral orbitofrontal regions.

**Fig. 4.**
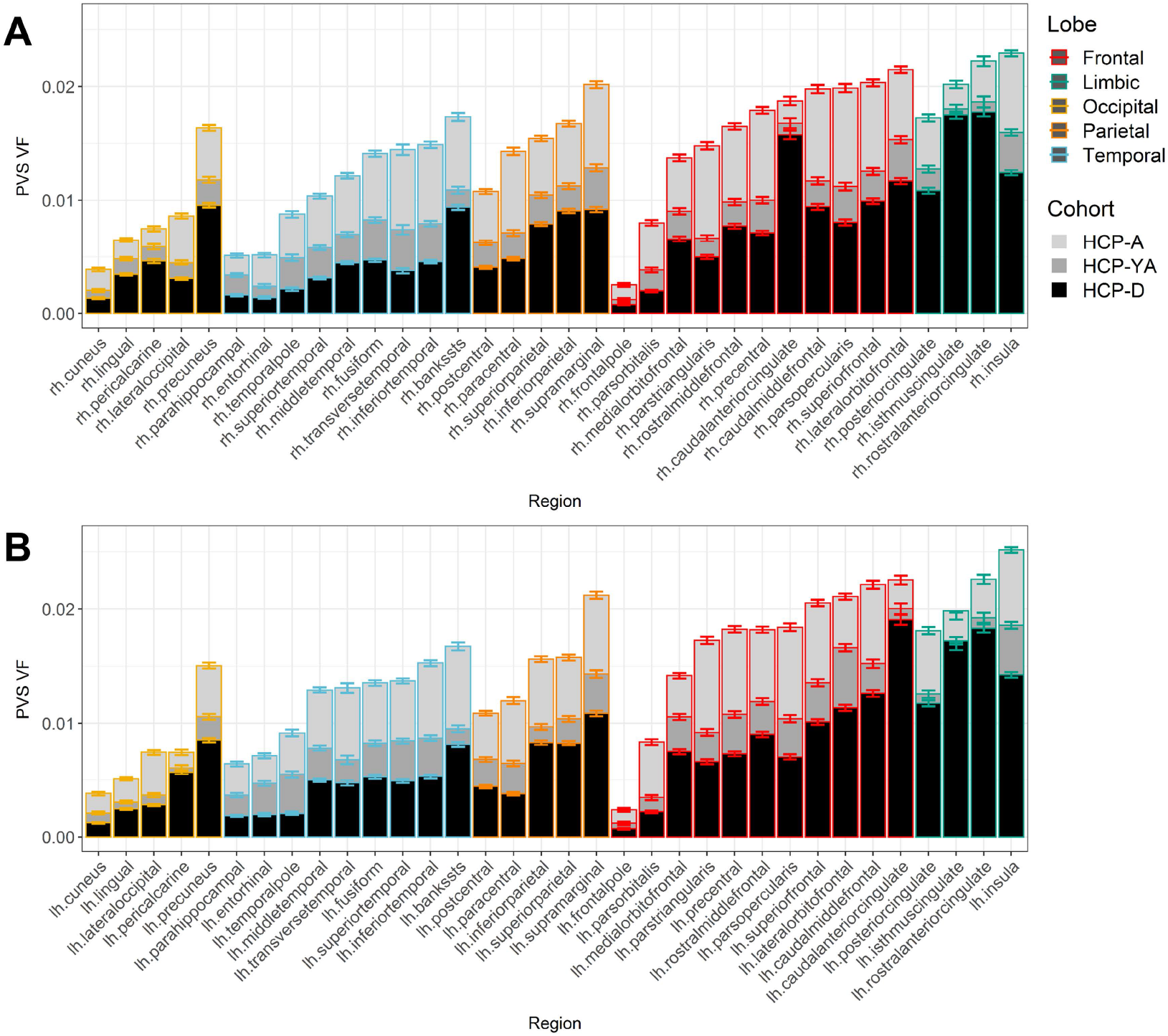
PVS VF varies across white matter regions. The mean PVS volume fraction across white matter regions in the (A) left and (B) right hemispheres are stratified by HCP cohort and ordered according to increasing PVS VF in the HCP-A cohort within major lobes. Occipital regions have the lowest PVS VF, while limbic and frontal regions have the highest. With some exceptions, regions with low PVS burden in aging subjects also tend to be low in young adults and developing subjects, while regions with high PVS burden in aging subjects tend to also be high in young adults and developing subjects. Bars represent standard error.

Exponential growth curves were used to quantify regional patterns of age-related PVS VF trajectories (**Fig. 5**). We found regions with the highest PVS VF in childhood are characterized by the slowest growth rate, including the white matter underlying the caudal anterior, isthmus and rostral anterior cingulate cortices bilaterally (**Fig. 5A-B**). Conversely, regions with low PVS VF in childhood, including white matter adjacent to bilateral cuneus, lateral and transverse occipital, inferior frontal and transverse temporal cortices had the largest growth rates. Indeed, we found the estimated PVS VF at age 8 years of age was significantly associated with slower growth rates across the lifespan (**Fig. 5C**; *B*=-.71, *t*(66)=-12.15, *p*<.001, *R^2^*=.69), where limbic regions tend to have high PVS VF in childhood that changes minimally with age and temporal regions have low PVS VF in childhood that show the largest percent difference across the lifespan (**Fig. 6**). Occipital regions deviate from this trend, where PVS VF remains low throughout the lifespan. These trends were observed bilaterally and the growth rate of left hemisphere white matter regions were significantly and positively correlated with the right hemisphere (**Fig. 5D**; *B*=.98, *t*(32)=16.43, *p*<.001, *R^2^*=.89).

**Fig. 5.**
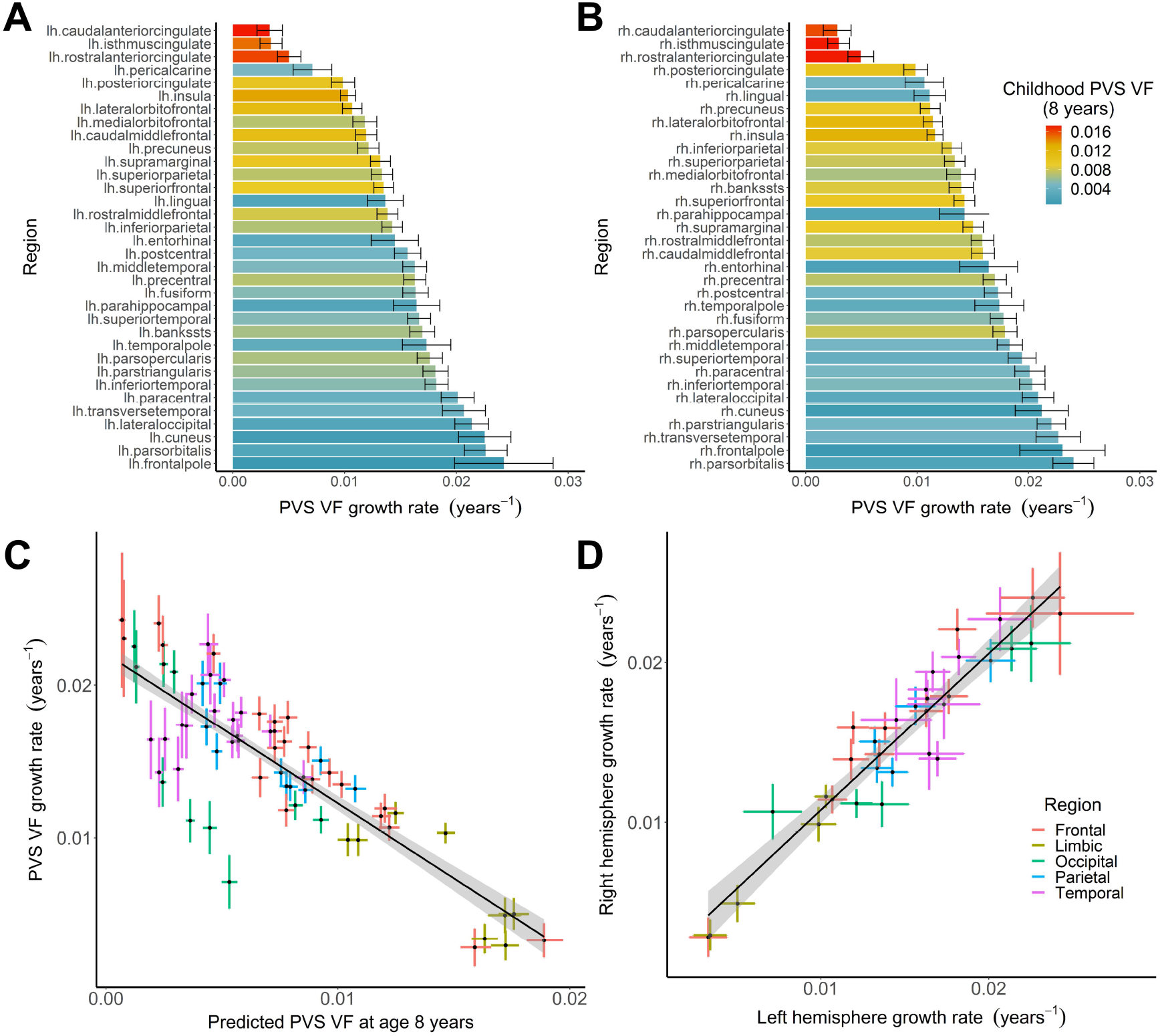
The PVS VF growth rate across the lifespan is driven by PVS burden in childhood. Age-related changes to PVS volume fraction were fit with an exponential model to assess regional differences in PVS lifespan trajectories. Bars denoting the growth rate, k, for age-related changes to the PVS volume fraction are shown for the (A) left and (B) right white matter regions ordered from lowest to highest. The color of the bars indicate the estimated PVS VF at age 8 years. (C) The relationship between PVS VF predicted at 8 years of age (x-axis) and the PVS growth rate (y-axis) across the lifespan for bilateral white matter regions is shown. Each point reflects a single white matter region with the 95% confidence interval estimated with bootstrap resampling. PVS burden is inversely correlated with the PVS VF growth rate across the lifespan, with trends in major regional demarcations. Limbic regions with high PVS burden in childhood undergo minimal growth across the lifespan, while temporal regions have low PVS burden in childhood and undergo rapid growth across the lifespan. Occipital regions are an exception to this finding, which tend to have low PVS burden in childhood and undergo minimal growth across the lifespan. (D) The estimated growth rate for PVS VF in white matter regions are highly correlated between the left and right hemispheres.

**Fig. 6.**
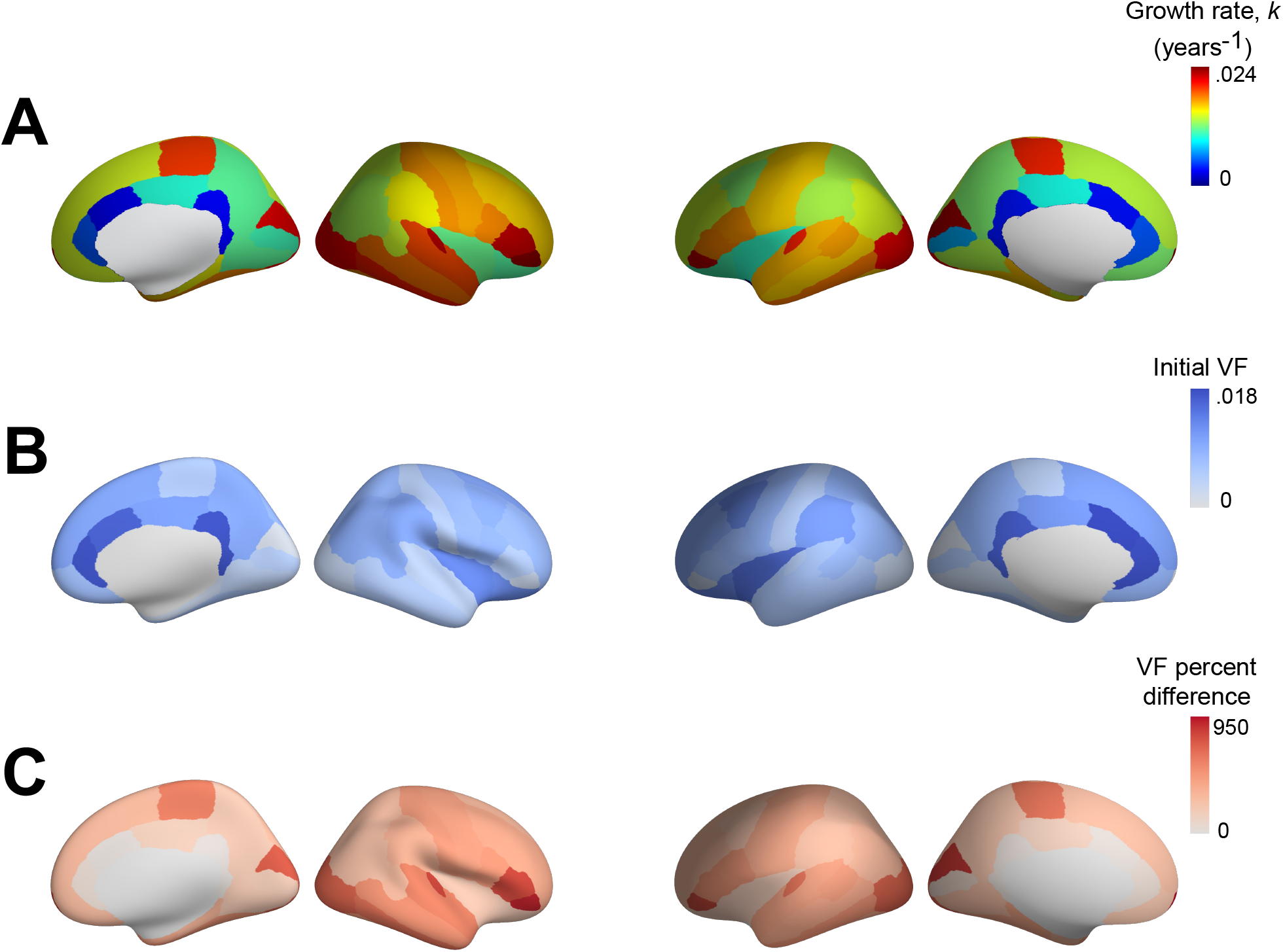
Regional patterns of PVS VF trajectories across the lifespan. The (A) growth rate, (B) PVS VF at age 8 years and (C) percent change in PVS VF from age 8 to 89 years are shown for the 34 bilateral white matter regions projected onto the corresponding Freesurfer cortical surface label.

### PVS sex differences

In the combined lifespan dataset, a main effect of sex on PVS VF (*F*(1,1381)=76.73, *p*<.0001), PVS count (*F*(1,1381)=106.31, *p*<.0001), and mean PVS solidity (*F*(1,1381)=77.85, *p*<.0001) were observed in the BG after controlling for age, scanner type and BG volume, where males had significantly larger PVS VF and count and smaller mean PVS solidity compared to females. The main effect of sex on mean PVS diameter in the BG did not survive multiple comparison correction (*F*(1,1381)=7.27, *p*=.007). Within white matter, a significant effect of sex on mean PVS solidity (*F*(1,1381)=63.83, *p*<.0001) and PVS VF (*F*(1,1381)=24.51, *p*<.0001) was observed, where males had larger PVS VF and smaller mean PVS solidity compared to females after controlling for age, scanner type and total white matter volume. The main effect of sex on white matter PVS count (*F*(1,1381)=5.36, *p*=.02), and mean diameter (*F*(1,1381)=4.64, *p*=.03) did not survive multiple comparison correction. The main effect of sex on PVS morphological features for each cohort are shown in **Table 4**.

**Table 4.** Sex differences in PVS morphological features stratified by HCP cohort

| Feature | Cohort | White matter |  |  | BG |  |  |
| --- | --- | --- | --- | --- | --- | --- | --- |
|  |  | *t* | Females | Males | *t* | Females | Males |
|  |  |  | Mean (SD) | Mean (SD) |  | Mean (SD) | Mean (SD) |
| VF | HCPD | 6.67*** | 6.25 x 10<sup>-3</sup> (2.87 x 10<sup>-3</sup>) | 8.16 x 10<sup>-3</sup> (3.29 x 10<sup>-3</sup>) | 6.33*** | 1.05 x 10<sup>-2</sup> (4.89 x 10<sup>-3</sup>) | 1.34 x 10<sup>-2</sup> (5.14 x 10<sup>-3</sup>) |
|  | HCPYA | 2.98** | 9.68 x 10<sup>-3</sup> (3.51 x 10<sup>-3</sup>) | 1.05 x 10<sup>-2</sup> (3.87 x 10<sup>-3</sup>) | -1.08 | 1.57 x 10<sup>-2</sup> (4.56 x 10<sup>-3</sup>) | 1.51 x 10<sup>-2</sup> (5.43 x 10<sup>-3</sup>) |
|  | HCPA | 0.08 | 1.55 x 10<sup>-2</sup> (4.66 x 10<sup>-3</sup>) | 1.57 x 10<sup>-2</sup> (4.42 x 10<sup>-3</sup>) | 7.08*** | 1.74 x 10<sup>-2</sup> (8.03 x 10<sup>-3</sup>) | 2.24 x 10<sup>-2</sup> (9.47 x 10<sup>-3</sup>) |
| Count | HCPD | 2.06* | 1101 (404) | 1374 (441) | 4.16*** | 181 (69) | 245 (68) |
|  | HCPYA | -1.37 | 1489 (431) | 1411 (382) | -1.06 | 209 (43) | 227 (64) |
|  | HCPA | 1.67 | 1713 (396) | 2045 (480) | 5.51*** | 177 (50) | 235 (62) |
| Diameter<br>(mm) | HCPD | 3.68*** | 1.79 (0.39) | 1.93 (0.42) | 0.69 | 1.31 (0.23) | 1.33 (0.23) |
|  | HCPYA | 3.31** | 1.92 (0.38) | 2.04 (0.43) | -0.63 | 1.45 (0.24) | 1.43 (0.22) |
|  | HCPA | -1.96 | 2.39 (0.45) | 2.33 (0.41) | 3.20** | 1.64 (0.28) | 1.71 (0.29) |
| Solidity | HCPD | -4.43*** | 0.61 (0.03) | 0.60 (0.03) | -4.54*** | 0.58 (0.03) | 0.57 (0.03) |
|  | HCPYA | 1.23 | 0.59 (0.03) | 0.59 (0.03) | -2.59** | 0.56 (0.03) | 0.56 (0.03) |
|  | HCPA | -3.17** | 0.57 (0.03) | 0.57 (0.03) | -6.32*** | 0.56 (0.03) | 0.54 (0.04) |

Main effect of sex on PVS morphological features in subcortical white matter and BG per HCP cohort after controlling for the effects of sex. PVS count is additionally normalized by intra-cranial volume.

While PVS burden in the BG was consistently larger in males compared to females across the lifespan (***SI Appendix*, Fig. S2**), differences in the age-related trajectories of PVS morphological alterations were observed between sexes in the white matter (***SI Appendix*, Fig. S3**). After controlling for covariates, a significant interaction between age and sex on white matter PVS VF (*F*(3,1381)=17.62, *p*<.0001) was observed, such that the covariate-adjusted rate of increase in PVS VF was greater in females (*β*=2.13×10^−4^, *t*(782)=34.30, *p*<.0001, adjusted *R^2^*=.60), compared to males (*β*=1.75×10^−4^, *t*(603)=24.11, *p*<.0001, adjusted *R^2^*=.49). Similarly, an interaction between age and sex on PVS diameter was also observed (*F*(3,1381)=21.95, *p*<.0001), where the adjusted mean diameter increased faster with age in females (*β*=.014, *t*(782)=20.53, *p*<.0001, adjusted *R^2^*=.34), compared to males (*β*=.009, *t*(603)=11.14, *p*<.0001, adjusted *R^2^*=.17). The cohort-stratified analyses also show males have significantly greater PVS VF and larger mean diameters compared to females in HCP-D and HCP-YA, while no significant sex differences were observed in HCP-A (***SI Appendix*, Table S1**). No significant interactions between age and sex were observed for the remaining PVS features.

### PVS associations with vascular risk factors

The HCP-YA and HCP-A cohorts were combined into a single dataset (n=918) to assess the influence of vascular risk factors on PVS enlargement patterns across the lifespan after controlling for age, sex and scanner type in the analyses. BMI showed the strongest associations with PVS morphology in the white matter (***SI Appendix*, Figure S4**)

Within the white matter, BMI was significantly associated with larger PVS VF (*F*(7,911)=36.02, *p*<.0001) and count (*F*(7,911)=73.11, *p*<.0001) and reduced mean diameter (*F*(7,911)=13.22, *p*=.0003) and solidity (*F*(7,911)=102.78, *p*<.0001) after controlling for age, sex and scanner environment. A significant interaction between BMI and age was observed for PVS diameter in the white matter (*F*(8,910)=6.83, *p*=.009), where PVS diameter increased faster with age in healthy weight participants (BMI<25) compared to overweight and obese participants (BMI>25). Within the BG, no significant relationship between PVS morphology and BMI was observed.

Elevated blood pressure (>130/80 mmHg) was significantly associated with increased PVS VF in the white matter (*F*(7,674)=14.85, *p*=.0001). No morphological features in the BG were significantly associated with blood pressure status. The full model that consists of age, sex, BMI, scanner type, systolic blood pressure and diastolic blood pressure accounts for 46% of the variance in PVS VF (*F*(8,909)=96.27, *p*<.001). BMI explained 1.6% of the unique variance in PVS VF, while systolic and diastolic blood pressure account for .3% and .02% of the variance, respectively.

### PVS morphological associations

Participants with increased PVS burden in the white matter also tended to have increased PVS burden in the BG (**Fig. 7**). PVS morphological features were also significantly correlated with one another within the BG (***SI Appendix*, Fig. S5**) and subcortical white matter (***SI Appendix*, Fig. S6**), where PVS VF was associated with increased mean diameter and count and decreased mean solidity. In order to better understand the contribution of PVS morphological features to alterations in the overall volume of the PVS, multiple linear regression was carried out on standardized variables to predict the PVS VF from PVS count, diameter and solidity. Within the BG, PVS count, mean diameter and mean solidity collectively account for 82% of the variance in PVS VF (*F*(3,1385)=2098, *p*<.0001, adjusted *R^2^*=.819). Mean PVS diameter has the highest predictive power (*β*=.549, *t*(1385)=35.49, *p*<.0001), followed by PVS count (*β*=.449, *t*(1385)=31.76, *p*<.0001), and mean PVS solidity (*β*=.128, *t*(1385)=7.06, *p*<.0001). PVS mean diameter and PVS count explain 16% and 13% of the unique variance in PVS VF, respectively, while mean PVS solidity explains 1% of the unique variance in PVS VF. Relative importance analysis was used to understand the extent to which each PVS morphological feature drives the prediction of PVS volume in the multiple regression model. Mean PVS diameter contributes .327 (95% CI = [.295, .358]) to the total proportion of variance, while PVS count and mean solidity contribute .251 [.230, .276] and .241 [.225, .257], respectively.

**Fig. 7.**
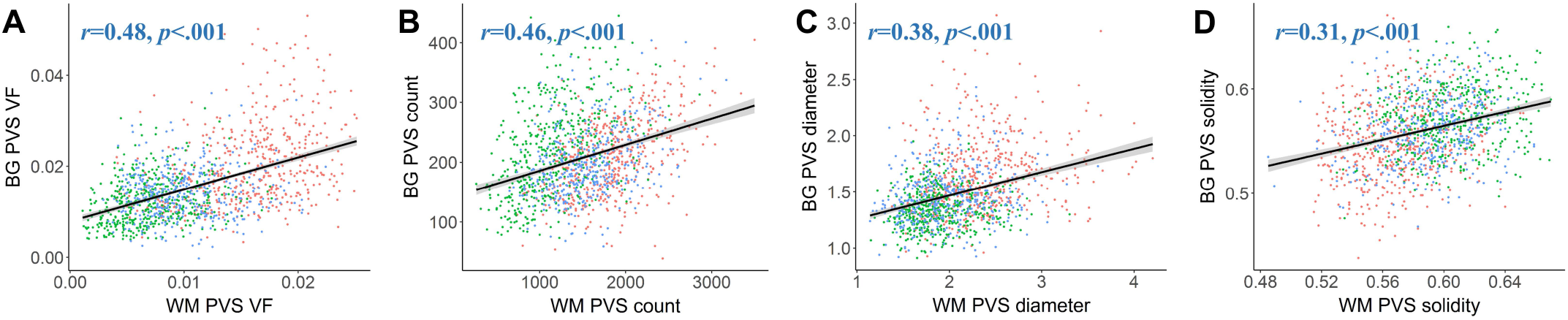
Similar distributions of PVS morphological features are found in the BG and white matter. The relationship between PVS morphology between the white matter (x-axis) and BG (y-axis) are shown for (A) PVS VF, (B) PVS count, (C) mean PVS diameter and (D) mean PVS solidity are shown. A significant and positive relationship between PVS features in the white matter and basal ganglia were found for all PVS features, where subjects with low PVS burden in the white matter also tended to have low PVS burden in the BG, and vice versa. Each point corresponds to the measurement of a single subject and is color-coded according to the HCP cohort: green – HCP-D, blue – HCP-YA, red – HCP-A

Within the white matter, PVS count, mean diameter and mean solidity account for 84% of the variance in PVS VF (*F*(3,1385)=2424, *p*<.0001, adjusted *R^2^*=.839). Mean PVS diameter has the highest standardized beta in multiple regression (*β*=.498, *t*(1385)=42.76, *p*<.0001), followed by count (*β*=.377, *t*(1385)=19.21, *p*<.0001), and mean solidity (*β*=.285, *t*(1385)=15.20, *p*<.0001). Mean PVS diameter explains 21.12% of the unique variance in PVS VF, while PVS count and mean solidity account for 4% and 3%, respectively. Relative importance analysis shows mean PVS diameter contributes .315 (95% CI = [.288, .340]) to the total proportion of variance, while PVS count and mean PVS solidity contribute .301 [.284, .319] and .224 [.206, .240], respectively.

## Discussion

We utilized an automated processing approach to automatically delineate and quantify PVS morphology in the BG and white matter to model PVS enlargement trajectories over an 80-year period of the lifespan. PVS visibility increases with age and is characterized by increased PVS volume fraction, count and caliber and decreased solidity. Furthermore, variability in PVS enlargement patterns is partially explained by anatomical location, sex and BMI. These findings are corroborated by previous studies that have found PVS visibility is apparent in children and adolescents (Groeschel et al., 2006; Piantino et al., 2020; Rollins et al., 1993) and increases with age in cognitively normal young adults (Barisano et al., 2020; Choi et al., 2020) and elderly subjects (Francis et al., 2019; Gutierrez et al., 2013; Huang et al., 2021; A. Laveskog et al., 2018; Yakushiji et al., 2014; Zhu et al., 2011, 2010). The present study builds upon these previous studies by considering PVS structure using multiple morphological characteristics across a broad age range in order to more specifically describe the time course of PVS alterations in healthy aging. Our finding of increased PVS count and caliber across the lifespan suggests the observed expansion of the PVS VF in previous studies is likely attributed to widening of the PVS, and not due to decreased tissue volume associated with age-related brain atrophy.

Increased visibility of PVS has been associated with a multitude of neurological conditions; however, it is unclear if PVS enlargement as a consequence of normal aging reflects neuropathology. Several factors may contribute to age-related enlargement of PVS, including physical alterations that disrupt bulk flow, such as changes to CSF pressure and peri-arterial blockages (Jessen et al., 2015). Previous studies have also shown normal aging is associated with impairments to the paravascular clearance pathway (Kress et al., 2014) due to age-related alterations in CSF production (Fleischman et al., 2012), arterial pulsatility (Zieman et al., 2005) and blood-brain barrier permeability (Farrall and Wardlaw, 2009). Therefore, it is hypothesized that age-related PVS enlargement may result in reduced fluid exchange and elimination from the brain indicative of dysfunctional waste clearance (Jessen et al., 2015; Mestre et al., 2017; Troili et al., 2020; Wardlaw et al., 2020). Because the highest risk factor for the development of neurodegenerative diseases is aging, such alterations to waste clearance mechanisms are speculated to contribute to the accumulation of protein aggregates and thus render the brain more vulnerable to neurodegenerative pathology or escalate the progression of pre-existing cognitive deficits (Jessen et al., 2015; Sweeney et al., 2018).

While PVS alterations were observed globally across the lifespan, distinct morphological trajectories were observed in the PVS of the BG and white matter. Age-related increases to PVS VF in the BG was predominantly observed in older adults within the HCP-A cohort, which coincided with wider diameters and, to a lesser extent, a larger number of visible PVS. Conversely, PVS VF in white matter increased linearly with age across the lifespan and was mainly attributed to increased PVS caliber, though PVS count increased up until middle age. We suspect the differing trajectories of PVS expansion may be attributed to differences in the anatomical characteristics that define vasculature in these structures, such as the cerebral arteries that feed into the network of perforating blood vessels that the PVS surround. BG PVS surround lenticulostriate arteries that branch from the M1 segment of the middle cerebral artery (MCA) (Marinković et al., 2001), which is the most pathologically affected vascular territory in the brain (Ng et al., 2007; Zaidat et al., 2014). There is converging evidence from several studies that show vascular pathologies are selectively associated with PVS enlargement in the BG, but not white matter (Charidimou et al., 2017, 2013; Doubal et al., 2010; Martinez-Ramirez et al., 2013; Potter et al., 2015). Because the risk for cerebrovascular disease increases with advancing age (Hauer et al., 2017), it is possible that the accelerated BG PVS enlargement observed in older subjects may be due, in part, to the increased incidence of pathological vascular influences in the aging cohort; however, we do not have sufficient information regarding vascular risk factors in this cohort to confirm this.

Conversely, white matter PVS envelop arteries that originate from multiple major cerebral arteries. The anterior cerebral artery (ACA) supplies blood vessels to anterior-frontal regions around the interhemispheric fissure, the MCA supplies parietal, temporal and inferior frontal regions via the M4 cortical segment and the posterior cerebral artery (PCA) supplies occipital regions (Kim et al., 2019). PVS were not uniformly distributed across the white matter, as we found frontal, parietal and cingulate regions had the highest PVS burden, while occipital and temporal regions have the lowest PVS burden bilaterally. Our findings are in line with previous studies that show the highest PVS burden in frontal and parietal lobes across all age groups (Barisano et al., 2020; Bouvy et al., 2014; A. Laveskog et al., 2018; Piantino et al., 2020). Additionally, a recent study in young adults found PVS VF in regions supplied by the ACA and MCA is elevated compared to those supplied by the PCA (Zong et al., 2020). Overall, we observed regionally-dependent patterns of PVS enlargement trajectories across the lifespan, where PVS levels in childhood were inversely correlated with the rate of growth. Regions fed by the ACA, such as cingulate and prefrontal white matter, have elevated PVS VF in childhood that remains relatively constant across the lifespan, while temporal, inferior frontal and lateral occipital regions supplied by the MCA have low PVS VF in childhood and undergo rapid enlargement across the lifespan. Medial occipital regions supplied by the PCA are an exception to this trend, as PVS VF levels remain low across the lifespan. Together, our findings suggest the mechanisms that contribute to age-related PVS enlargement differ according to the cerebral arteries that support each region.

It is unclear if the age-related increase in PVS cross-section diameter reflects pathological enlargement. From childhood to advancing age, the mean PVS diameter in the present study ranges from 1.85 mm to 2.36 mm in the white matter and 1.32 mm to 1.66 mm in the basal ganglia. Previous MRI studies acquired at 3T have similarly found the majority of PVS have a maximum diameter of 3 mm (Del C. Valdes Hernandez et al., 2013; Troili et al., 2020); however, it is likely that many more PVS go undetected in the brain due to limitations in the spatial resolution afforded by clinical MRI scanners. PVS with sub-millimeter diameters have been observed with ultra-high field MRI (Barisano et al., 2021), which provides significantly improved visualization of small PVS, and histological studies in vascular tissue suggest the majority of PVS diameters range from .13 to .96 mm (Pesce and Carli, 1988). Due to the preponderance of small-caliber PVS in the brain that are undetectable at the current resolution, it is feasible that the observed PVS diameter enlargement may be attributed to the simultaneous expansion of multiple neighboring PVS, thus resulting in the appearance of a single larger PVS (Barisano et al., 2021).

Our finding of increased PVS count with age corroborates previous studies (Bouvy et al., 2014; Zong et al., 2020) and is likely attributed to increased visibility of pre-existing PVS, as opposed to the formation of new ones, because PVS spatially localize with penetrating arterioles and venules that remain static across age. Consistent with our findings of a strong correlation between PVS count and mean diameter, increased PVS count could therefore reflect increased caliber of small PVS previously undetectable with MRI. The age-related increase in PVS count may also be attributed to discontinuities in PVS visibility, where a single PVS is delineated as multiple disconnected segments. While discontinuities may be attributed to partial volume effects in small PVS, others have suggested PVS enlargement can manifest as focal dilations (Wardlaw et al., 2013) that may be visualized as several separate segments. This latter interpretation is supported by our finding of age-related reductions in PVS solidity, a measure that describes shape complexity based on the fraction of voxels enclosed within the smallest complex polygon, where low solidity tends to describe highly tortuous paths (Westrate et al., 2014). Previous studies have similarly found increased arterial tortuosity with aging (Cha et al., 2003; Del Corso et al., 1998; Schnerr et al., 2017). It is unclear whether the increased complexity of the PVS observed in the present study is due to alterations to the PVS itself, through focal dilations and degradation, or if it is guided by the underlying structure of the vasculature. Together, our finding of increased PVS VF, count and diameter accompanied by decreased solidity across the lifespan point towards age-related enlargement and possibly degradation of PVS.

Inter-subject variability in PVS morphology is partially explained by sex and body mass, but not blood pressure. Previous studies have shown PVS volumes are larger in males compared to females in adult (Barisano et al., 2020; Zhu et al., 2010) and aging subjects (Ramirez et al., 2015). While we found PVS VF was significantly larger in males compared to females in the white matter and BG, males also had significantly more PVS compared to females in the BG, specifically. Androgen and estrogen receptors on vascular endothelial cells act as direct targets for circulating sex hormones (Zuloaga et al., 2012b, 2012a) and the relative distributions of these hormones across the lifespan may contribute to the observed PVS sex differences. While androgens have been shown to protect the blood-brain barrier (Atallah et al., 2017), they can also adversely affect vasculature through neuroinflammatory stimulation (Gonzales et al., 2009) and vasoconstriction (Gonzales et al., 2004). Therefore, the unique actions of circulating androgens on the cerebrovascular system may further exacerbate age-related PVS enlargement in males. We also found age-related increases to the volume and diameter of white matter PVS occur more rapidly in females compared to males, where PVS burden in females is lower in childhood and adulthood and equalizes with males in older subjects. Estrogen is atheroprotective and has been shown to suppress the cerebrovascular inflammatory response (Razmara et al., 2008; Sunday et al., 2006), reduce blood-brain barrier permeability (Burek et al., 2010) and induce vasodilation (Murata et al., 2013). However, the protective effects of estradiol wanes with age as the gonadal production of sex hormones declines after menopause (Wildman et al., 2008) and is reflected by an age-related increase in blood-brain barrier permeability in females compared to males (Liu et al., 2009). Therefore, the accelerated PVS enlargement in females compared to males across the lifespan may be attributed to age-related reductions in circulating estrogen levels.

Within the white matter, excess body mass was associated with increased PVS volume and count, but decreased diameter and slower enlargement patterns compared to normal weigh adults. These seemingly contradictory findings of increased PVS burden and decreased PVS size suggests excess body mass is associated with increased visibility of predominantly small diameter PVS. It is possible that the enlarged PVS observed in healthy weight adults may not represent vascular pathology but may instead reflect a phenomenon of normal aging. Our lack of a significant relationship between BMI and PVS morphology in the BG is surprising, given previous research that has shown both BMI (Arnoldussen et al., 2019) and lenticulostriate PVS (Francis et al., 2019) are tightly coupled with cerebrovascular risk factors. However, elevated BMI also promotes systemic inflammation, and previous studies have shown white matter PVS pathology in neuroinflammatory disease, such as multiple sclerosis. The variance in age-related PVS alterations was not adequately explained by blood pressure differences. There is strong evidence to link PVS size with hypertension, particularly within the BG; however, only 61 participants in the present study had high blood pressure (>140/90 mmHg) and it is likely that our study did not have sufficient subjects with clinical hypertension to detect PVS alterations related to high blood pressure. Therefore, our findings suggest that sub-clinical differences in blood pressure may not significantly contribute to PVS morphological variability.

While the current study utilizes a granular approach to explore the spatial distribution of PVS morphological characteristics across age, several limitations should be considered. HCP is a vast dataset acquired on different scanners at multiple sites and data for the developing and aging cohorts were acquired with slightly different MRI acquisitions than the young adult cohort. To overcome this, we employed robust post-processing harmonization techniques to account for differences in acquisitions and controlled for scanner type as a covariate of no interest in our analyses. Furthermore, we analyzed cohorts separately and found these results were consistent with our lifespan trends. Another limitation was the cross-sectional nature of our study and associations between PVS morphology and age point towards differences, not within-subject changes, across the lifespan. Our research is also limited by the spatial resolution afforded with clinically feasible MRI. In a recent simulation study, researchers found structural MRI data can accurately quantify PVS with diameters greater than twice the image resolution (Bouvy et al., 2014). Therefore, the presence of small PVS may lead to a systematic over-estimation of PVS diameters.

The present study sought to characterize the time course of PVS morphological alterations in the BG and subcortical white matter regions across the lifespan in a large cross-sectional cohort of ∼1400 cognitively normal subjects. We found PVS within white matter regions and the BG undergo distinct age-related trajectories that may be indicative of their respective roles in pathological conditions characterized by failure of the waste clearance system. Furthermore, the rate of PVS enlargement in white matter regions were dependent on the degree of PVS burden in childhood, highlighting a critical link between perivascular physiology in childhood and in aging. The findings presented here will aid in our understanding of the role of PVS in health and disease by providing a benchmark to which pathological PVS alterations can be compared. Therefore, future studies should aim to understand the pathophysiological and normative mechanisms that give rise to PVS enlargement in order to better assess its utility as a diagnostic biomarker for neurodegenerative and cerebrovascular disease.

## Materials and Methods

### Study Participants

Cognitively normal subjects from childhood through advanced age were recruited and scanned through the Washington University-University of Minnesota (WU-Minn) Human Connectome Project and the Lifespan Human Connectome Project in Development and Aging. Structural neuroimaging and demographic information from 1389 participants (784 females) across the lifespan were included in the present study (*M*±*SD*=34.17±20.07 years, 8.08 – 89.75 years) (**Table 1**).

#### HCP Development (HCP-D)

The sample of typically developing children and adolescents used in the present study were derived from the Lifespan Human Connectome Project in Development (Somerville et al., 2018). Subjects were recruited and scanned in Boston, Los Angeles, Minneapolis and St. Louis. Participants were excluded if they were born premature, require special educational services, had MRI contraindications, or had a history of serious medical problems, head injury, endocrine disorders, psychiatric disorders, or neurodevelopmental disorders. The goal of the HCP-D was to enroll at least 1,300 participants; however, 655 subjects (332 female) with neuroimaging data were available at the time of this study. Of the participants originally considered, 140 subjects were excluded in the HCP-D 2.0 release of the data, 32 were excluded due to poor structural MRI data quality, 6 subjects were excluded due to processing pipeline failures, and 1 subject was excluded due to anatomical abnormalities, resulting in the inclusion of 471 subjects (260 female) from the HCP-D cohort in the present analyses (*M*±*SD*=14.36±3.77 years, 8.08 – 21.92 years).

#### HCP Young Adults (HCP-YA)

The sample of healthy young adults were acquired through the WU-Minn Human Connectome Project at Washington University in St. Louis (Van Essen et al., 2013). HCP-YA utilized the same exclusionary criteria as the HCP-D; HCP-YA also excluded subjects with scores below 25 on the mini-mental state exam (MMSE). The study aimed to recruit 1,200 subjects; however, 1112 subjects with neuroimaging data were available at the time of the present study. Of those considered, 51 subjects were excluded due to poor structural data quality, 643 subjects were randomly excluded due to familial relationships with other participants (siblings), 12 subjects were excluded due to processing pipeline failures, and 1 subject was excluded due to anatomical abnormalities, resulting in a total of 405 subjects from the HCP-YA cohort (226 females) used in the present analyses (*M*±*SD*=28.73±3.78 years, 22.00 – 37.00 years).

#### HCP Aging (HCP-A)

The sample of cognitively normal aging adults older than 36 years of age were acquired through the Lifespan Human Connectome Project in Aging (Bookheimer et al., 2019). Participants were recruited and scanned at Washington University St Louis, University of Minnesota, Massachusetts General Hospital and the University of California Los Angeles. In addition to the exclusionary criteria listed for HCP-D and HCP-YA, individuals with impaired cognitive abilities were excluded from HCP-A according to tiered age-appropriate cut-off scores. While over 1,200 subjects will ultimately be enrolled in the study, there were 687 subjects with neuroimaging data available at the time of analysis. Of those originally considered, 105 subjects were excluded following the 2.0 release of the HCP-A dataset, 49 were additionally excluded due to poor structural data quality following quality assurance purposes, 19 subjects were excluded due to processing pipeline failures, and 1 subject was excluded due to anatomical abnormalities, resulting in the inclusion of a total of 513 subjects (298 females) from the HCP-A cohort in the present analyses (*M*±*SD*=56.35±13.65 years, 36.00 – 89.75 years).

### MRI Acquisition

#### HCP-YA

Participants were scanned on a Siemens 3T Connectome Skyra with a 100 mT gradient coil and a 32-channel Siemens receive head coil (Glasser et al., 2013). B0 field maps, B1- and B1+ maps were collected to correct for readout distortion and intensity inhomogeneities. T1-weighted (T1w) images were collected as a 3D single-echo magnetization prepared – rapid gradient echo (MP-RAGE) images with the following acquisition parameters: voxel size=.7 mm isotropic, FOV = 224 mm, matrix = 320, 256 sagittal slices per slab, TR = 2400 ms, TE = 2.14 ms, TI = 1000 ms, FA = 8 degrees, Bandwidth = 210 Hz per pixel, echo spacing = 7.6 ms, GRAPPA factor = 2, 10% phase encoding oversampling (A-P), dwell time=7.4 μs. T2-weighted (T2w) images were collected as 2 variable flip angle turbo spin-echo sequences averaged together with the following acquisition parameters: voxel size=.7 mm isotropic, FOV = 224 mm, matrix = 320, 256 sagittal slices per slab, TR = 3200 ms, TE = 565 ms, BW = 744 Hz/pixel, no fat suppression pulse, GRAPPA = 2, turbo factor = 314, echo train length = 1105 echoes, 10% phase encoding oversampling, dwell time = 2.1 μs.

#### HCP-D and HCP-A

Slight variations in the acquisition parameters were made for the HCP-A and HCP-D cohorts to accommodate the unique challenges of working with young and elderly populations (Harms et al., 2018). HCP-D and HCP-A participants were scanned on a variant of the HCP-YA Connectome scanner, the Siemens 3T Prisma with an 80 mT/m gradient coil and a Siemens 32-channel Prisma head coil. T1w multi-echo MP-RAGE scans were acquired with the following acquisition parameters: voxel size=.8 mm isotropic, 4 echoes per line of k-space, FOV = 256 x 240 x 166 mm, matrix = 320 x 300 x 208 slices, 7.7% slice oversampling, GRAPPA = 2, pixel bandwidth = 744 Hz/pixel, TR = 2500 ms, TI = 1000 ms, TE = 1.8/3.6/5.4/7.2 ms, FA = 8 degrees. Motion-induced re-acquisition were allowed for up to 30 TRs. T2w turbo spin echo (TSE) scans were collected from each subject with the following acquisition parameters: voxel size=.8 mm isotropic, 4 echoes per line of k-space, FOV = 256 x 240 x 166 mm, matrix = 320 x 300 x 208 slices, 7.7% slice oversampling, GRAPPA = 2, pixel bandwidth = 744 Hz/pixel, TR = 3200 ms, TE = 564 ms, turbo factor = 314. Motion-induced re-acquisition were allowed up to 25 TRs.

### Quality assurance procedures

Prior to data processing and PVS segmentation, all T1w and T2w images were visually inspected and scans with poor data quality were excluded from analyses. Poor scan quality was characterized by T1w, T2w or EPC volumes with excessive head motion, reduced GM/WM tissue contrast, unidentifiable anatomy, blurred images and salt-and-pepper noise, which obstructed the visibility of PVS. Additionally, EPC images with contrast inhomogeneities due to slight variations in T1w and T2w data quality or registration failures were excluded.

### MRI data preprocessing

All T1w and T2w images were preprocessed using the HCP minimal processing pipeline version 4.0.1(Glasser et al., 2013) in parallel with the LONI pipeline (Dinov et al., 2009) . HCP-A and HCP-D structural images were re-sampled to a spatial resolution of .7 mm isotropic to match that of HCP-YA. Following gradient distortion correction, T2w images were aligned to T1w images using rigid body registration and then images were transformed to MNI space and brought into alignment with the anterior and posterior commissure (ACPC). Brain extraction and readout distortion correction using a field map were then carried out. T2w images were registered to subject T1w native space using boundary-based registration (BBR) with 6 degrees of freedom (Greve and Fischl, 2009). T1w and T2w images were corrected for readout distortion using the receive (B1-) and transmit (B1+) field maps and then registered to MNI space with an affine transformation implemented with FLIRT followed by a nonlinear registration implemented with FNIRT.

### Subcortical segmentation

In order to characterize regional subcortical PVS features, T1w images were processed with Freesurfer version 6 (http://surfer.nmr.mgh.harvard.edu/). Images were first downsampled to a voxel size of 1 mm isotropic to accommodate spatial resolution limitations of Freesurfer (Glasser et al., 2013). Data processing included motion correction, intensity normalization, removal of non-brain tissue and automated Talairach transformation (Reuter et al., 2010; Segonne et al., 2004; Sled et al., 1998). Subcortical white matter and BG were segmented using an atlas-based approach (Fischl et al., 2004, 2002). Subcortical white matter was further parcellated into 68 regional volumes (34 regions per hemisphere) labeled according to the nearest cortical label (Fischl et al., 2004, 2002) defined by the Desikan-Killiany atlas (Desikan et al., 2006).

### Perivascular space segmentation

PVS were automatically segmented and quantified using the processing technique described in(Sepehrband et al., 2019). Adaptive non-local mean filtering using image self-similarity properties was applied to the co-registered and bias field-corrected T1w and T2w images (Manjón et al., 2010). In order to increase visibility of the PVS in the subcortical white matter and BG for segmentation, an enhanced PVS contrast (EPC) was then generated by dividing the T1w image by the T2w image (Sepehrband et al., 2019).

Frangi filters (Frangi et al., 1998) were applied to the EPC using the Quantitative Imaging Toolkit (QIT) (Cabeen et al., 2018) to create PVS vesselness maps, which describes the extent to which a voxel corresponds to the tubular structure of the PVS. Frangi filter parameters were set to default values of *α* = *β* = 0.5 and *c* was set to half the maximum Hessian norm according to Frangi *et al*. (1998) (Frangi et al., 1998). The vesselness scale ranged from 0.1 to 5 voxels. Binary PVS maps were generated by thresholding the vesselness maps using a value optimized for the highest concordance and correlation with PVS counts from expert readers (scaled *t* = 1.5). The CSF volumetric mask generated by Freesurfer was dilated using a spherical kernel with a 1.4 mm radius and used to exclude voxels at the interface between white matter and ventricles to avoid incorrect classification of outside tissue voxels.

White matter hyperintensities (WMH) are anomalous structural features indicative of tissue pathology (Wardlaw et al., 2015). WMH is more common in the normal aging brain compared to younger brains (Zhuang et al., 2018) and has a similar contrast to PVS. While vesselness thresholding excludes most large globular WMH from consideration during PVS segmentation, incorrect classification of some small WMH as PVS remains a possibility. The influence of WMH on PVS segmentation was assessed in a subset of 184 participants (91 HCP-A, 50 HCP-YA, and 43 HCP-D subjects), where PVS maps were visually inspected and manually edited to remove segmentation false positives by a group of 4 trained raters, including WMH, ventricular walls due to inaccurate tissue segmentation and other erroneously labeled regions with hyper-intense EPC contrast. To ensure consistency among raters, 33% of the subjects were inspected by more than one rater. The corrected and uncorrected PVS maps were compared to determine if manual correction significantly improves analyses. The severity of WMHs in the EPC contrast was scored on a scale from 0 to 3 based on the number and size of WMH using the Fazekas scale (Prins and Scheltens, 2015), where 0 represents a scan with no identifiable WMH and 3 reflects several large WMHs. Out of the subjects randomly selected for analysis, 153 (83%) did not present with any observable WMH, while 11 (5.9%) were noted to have moderate (8 HCP-A subjects) or severe (3 HCP-A subjects) WMH. Overall, the false positive rate across all cohorts was 4.5% (HCP-D: 4.1%, HCP-YA: 7.3%, HCP-A: 3.1%). WMH severity was not significantly correlated with the PVS segmentation false positive rate (*r*(182)=.06, *p*=.40). Therefore, it is unlikely that PVS segmentation false positives due to increased WMH burden in older adults contributes to the observed age-related PVS alterations in the present study.

### Perivascular space feature extraction

The PVS volume fraction was defined as fraction of the volume of space occupied by PVS and was calculated by dividing the PVS volume by the subcortical white matter and basal ganglia volume. The PVS morphological properties of count, mean diameter and mean solidity were extracted from the 3-D volumetric mask of PVS using the Matlab function *regionprops3*. The PVS count summed up the number of contiguous PVS-labeled voxels within a given region of interest. PVS diameter reflects the average cross-sectional diameter and is calculated using the mean of the second and third principal eigenvectors of an ellipsoid constructed with the same normalized second central moments as the segmented PVS. The PVS solidity corresponds to the proportion of the PVS-labeled voxels within the convex hull that contains a given PVS segment. Because segmented PVS are not always cluster of voxels, The PVS mean solidity was calculated on PVS with a length of 5 contiguous voxels or greater. For analyses, the mean PVS solidity and PVS diameter are provided for each white matter and BG region of interest.

### Vascular risk factors

To assess the influence of vascular risk factors on PVS morphology, a portion of the analyses were performed in a subset of the HCP Lifespan that consisted of subjects older than 21 years of age in the HCP-YA and HCP-A cohorts (*n*=918).

Body mass index (BMI) was calculated from height (cm) and weight (kg) measurements collected during participant interviews using the equation: BMI = (weight)/(height)^2^. Participants were stratified into BMI groups according to World Health Organization (WHO) criteria for weight categories (“Obesity: preventing and managing the global epidemic. Report of a WHO consultation.,” 2000): (1) a “high” BMI group (*n*=529, 21.8 – 87.75 years, 45.2±17.2 years) that included subjects described as “overweight” or “obese” with BMI greater than or equal to 25 kg/cm^2^ (25– 45.17 kg/cm^2^, 29.8±4.0 kg/cm^2^) (and (2) a “low” BMI group (*n*=390, 21.8 – 89.8 years, 43.1±17.5 years) that includes subjects described as “healthy” with BMI less than 25 kg/cm^2^ (16.6– 24.9 kg/cm^2^, 22.4±1.8 kg/cm^2^).

Systolic and diastolic blood pressure were measured from participants in the seated position. To assess the influence of blood pressure on age-related PVS morphological alterations, subjects were stratified into groups: (1) A normal blood pressure group where systolic and diastolic blood pressure was less than 130 mmHg and 80 mmHg, respectively (*n*=297, 21.8-89.8 years, 39.2±14.2 years) and (2) An elevated blood pressure group where systolic and diastolic blood pressure was greater than or equal to 130 mmHg and 80 mmHg, respectively (n=384, 22.8-86.5 years, 46.2±16.9 years), per the American Heart Association guidelines (Whelton et al., 2018)

### Statistical analysis

In order to assess age-related alterations to PVS features (mean cross-sectional diameter, mean solidity, count and VF) across the lifespan, the HCP lifespan cohorts were analyzed both separately and combined as a single dataset. In an effort to adjust for non-biological sources of variance inherent in multi-site research studies, PVS morphometric features were first harmonized using ComBat-GAM (https://github.co m/rpomponio/neuroHarmonize). ComBat-GAM is an extension of the batch effect correction tool ComBat commonly used in genomics and recently developed to account for multi-site scanner effects. It has been previously shown that ComBat removes unwanted scanner effects, while preserving biological associations, in a large multi-study analysis that investigated cortical thickness in health and disease (Fortin et al., 2018). ComBat-GAM was run within an empirical Bayes framework, which assumes features share the same common distribution, and utilized a generalized additive model (GAM) with a penalized non-linear term to model the dynamic age effects typically observed across the lifespan (Pomponio et al., 2020). ComBat-GAM was applied to the PVS metrics extracted from the BG and white matter regions to correct for HCP cohort effects while preserving the effects of sex and the nonlinear effects of age.

Statistical analysis was then carried out with R version 3.1.2. In the combined dataset, the main effect of sex and age*sex interaction on harmonized PVS features after controlling for covariates was assessed with ANCOVA. For the main effect of age, Bayesian information criteria (BIC) was used to identify the model that accounts for the most age-related variance in PVS morphology among the following 4 models typically used to characterize lifespan trajectories: (1) linear regression, (2) second-order polynomial model, (3) Poisson curve, and (4) nonlinear growth curve. For the PVS features within the total white matter and BG, the best fit models were either linear regression (PVS = *B_0_* + *B_1_*\*age) or the second-order polynomial model (PVS = *B_0_* + *B_1_*\*age + *B_2_*\*age^2^). A nonlinear growth curve of the form PVS = *a*e*^k^*^*age^ was fit for each parcellated white matter region to explain the relationship between PVS VF and age in order to compare the rate of PVS VF change among regions, where *k* reflects the growth rate. The 95% confidence interval for each growth model coefficient and predicted age at 8 years of age were generated with bootstrap resampling with replacement (*N*=10,000). Statistical tests were corrected for multiple comparisons using a Bonferroni-adjusted significance threshold of p<.006 (.05/8).

Statistical tests for the main effects of age and sex on PVS features in the white matter and BG were also replicated separately within each HCP cohort. The following general linear model (stats v.3.6.2) was fit for each cohort separately to assess the influence of age on PVS structure: PVS = *B_0_* + *B_1_*\*age + *B_2_*\*sex + *B_3_*\*scanner + B*_4_*\*volume, where the dependent variable reflects the PVS morphological feature of interest and sex, scanner type and total tissue volume of either the white matter or BG are covariates of no interest. The main effect of sex after accounting for age, scanner type and tissue volume were assessed with analysis of covariance (ANCOVA). Statistical tests were corrected for multiple comparisons (3 cohorts x 4 PVS features x 2 structures) using a Bonferroni-adjusted significance threshold of p<.002 (.05/24).

## Data sharing plans

All data used in this study is available through the Human Connectome Project (HCP) (https://www.humanconnectome.org/). The HCP Young Adult 1200 Subject Release (S1200) can be downloaded at ConnectomeDB (https://db.humanconnectome.org/). The Lifespan HCP datasets (HCP Aging and HCP Development) can be accessed through the Connectome Coordination Facility (CCF) in the NIMH Data Archive (NDA) at https://nda.nih.gov/ccf (DO<u>I: 10.15154/1524651)</u>. Normative PVS data for brain regions across age groups will be made available through an online portal for data discovery.

## Acknowledgments

The image computing resources provided by the Laboratory of Neuro Imaging Resource (LONIR) at USC are supported in part by National Institutes of Health (NIH) National Institute of Biomedical Imaging and Bioengineering (NIBIB) grant P41EB015922. Author KML is supported by the National Institute on Aging (NIA) of the NIH Institutional Training Grant T32AG058507. The research reported in this publication was supported by the National Institute of Mental Health (NIMH) of the NIH under the award number RF1MH123223, the NIA of the NIH under the award number R01-AG070825, and the USC ADRC 1P30AG066530-01.

## Author Contributions

<u>Kirsten M. Lynch</u>: Conceptualization, Methodology, Formal analysis, Writing – Original Draft, Writing – Review and Editing, Visualization; <u>Farshid Sepehrband</u>: Conceptualization, Methodology, Software, Writing – Review & Editing, Resources, Data Curation, Supervision; <u>Arthur W. Toga</u>: Resources, Writing – Review & Editing, Supervision, Funding acquisition; <u>Jeiran Choupan</u>: Resources, Data Curation, Software, Writing – Review & Editing, Supervision, Project administration, Funding acquisition

## Competing Interest Statement

The perivascular space mapping technology is part of a pending patent owned by FS and JC, with no financial interest/conflict.

## Materials and Correspondence

Correspondence and requests for materials should be addressed to KML or JC

## Notes

https://db.humanconnectome.org/

https://nda.nih.gov/ccf

### Summary of Updates

Section on working memory performance has been removed; Included additional analyses on BMI and blood pressure to explain age-related PVS variance; Supplemental files updated

