## Supplementary material for "Brain perivascular space imaging across the human lifespan": SI Appendix, Figure S1

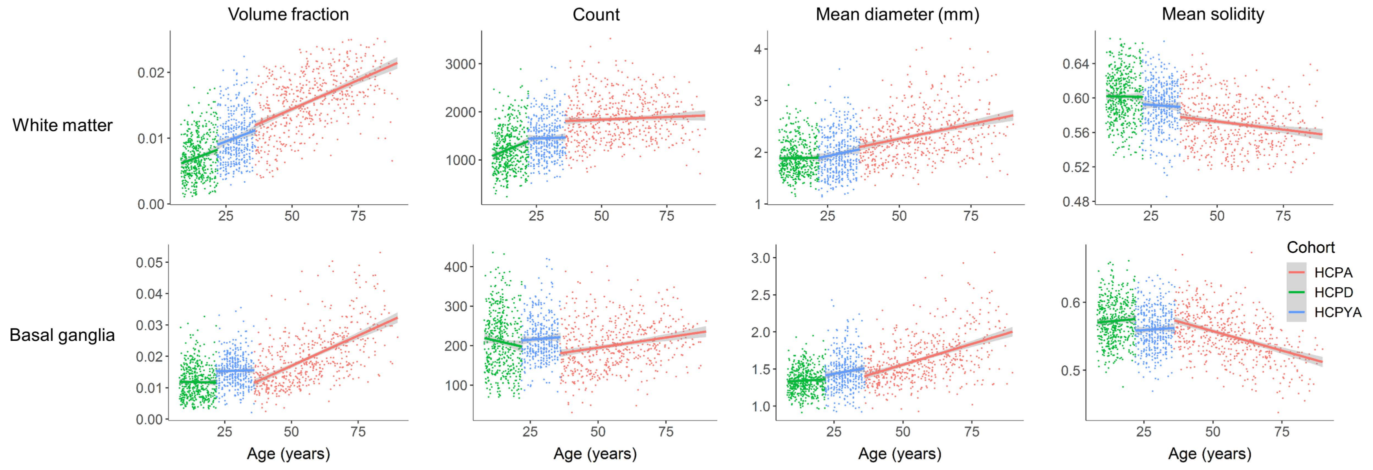


**Fig. S1.** Associations between PVS morphological features and age in the white matter (top) and basal ganglia (bottom) stratified by cohort. Linear regression lines of best fit are shown separately for each HCP cohort.
