## Supplementary material for "Brain perivascular space imaging across the human lifespan": SI Appendix, Figure S2

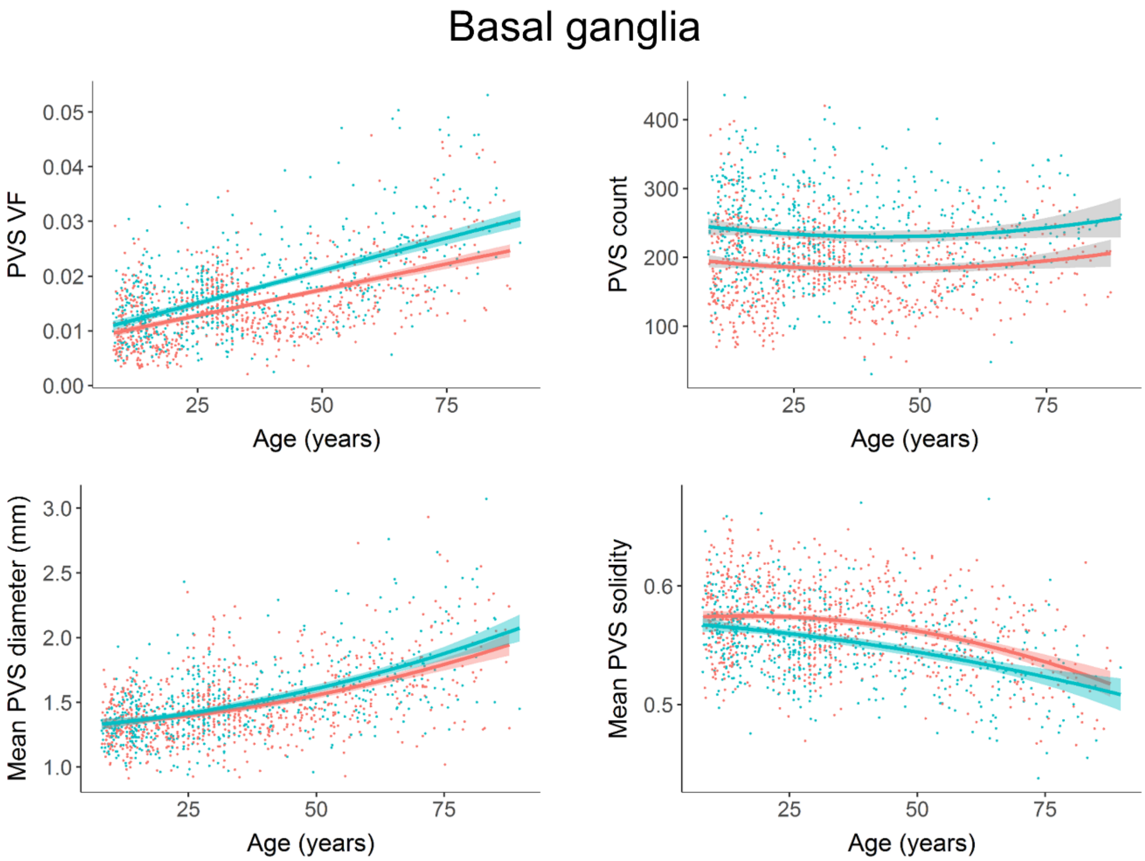


**Fig. S2.** Sex-stratified lifespan trajectories for PVS morphological features within the basal ganglia are shown where Males have more pronounced PVS burden across the lifespan in the basal ganglia. Individual data points, regression lines and shaded standard error estimates are colored according to sex, where blue reflects males and red denotes females.
