## Supplementary material for "Brain perivascular space imaging across the human lifespan": SI Appendix, Figure S3

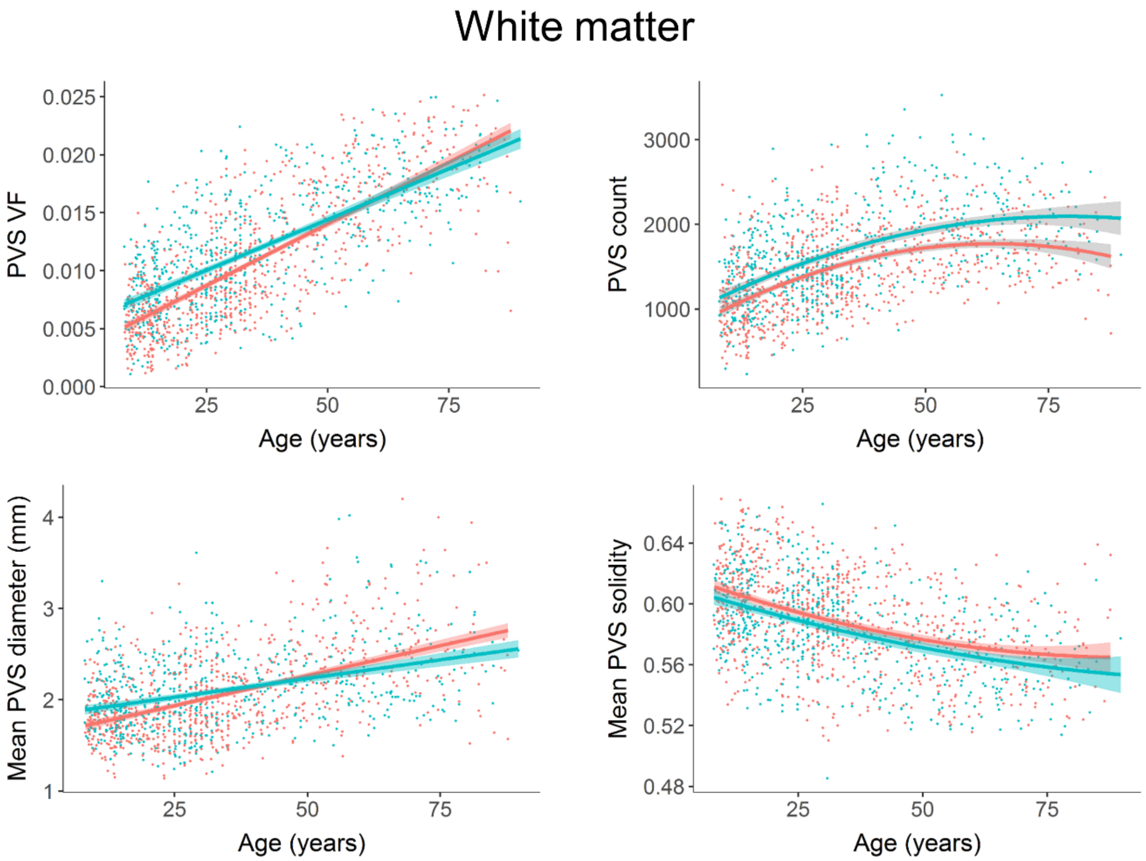


**Fig. S3.** Sex-stratified lifespan trajectories for PVS morphological features within the white matter are shown. Males and females have differing PVS VF and mean diameter trajectories across the lifespan in the white matter, where age-associated increases in features are more rapid in females compared to males. Individual data points, regression lines and shaded standard error estimates are colored according to sex, where blue reflects males and red denotes females.
