## Supplementary material for "Brain perivascular space imaging across the human lifespan": SI Appendix, Figure S4

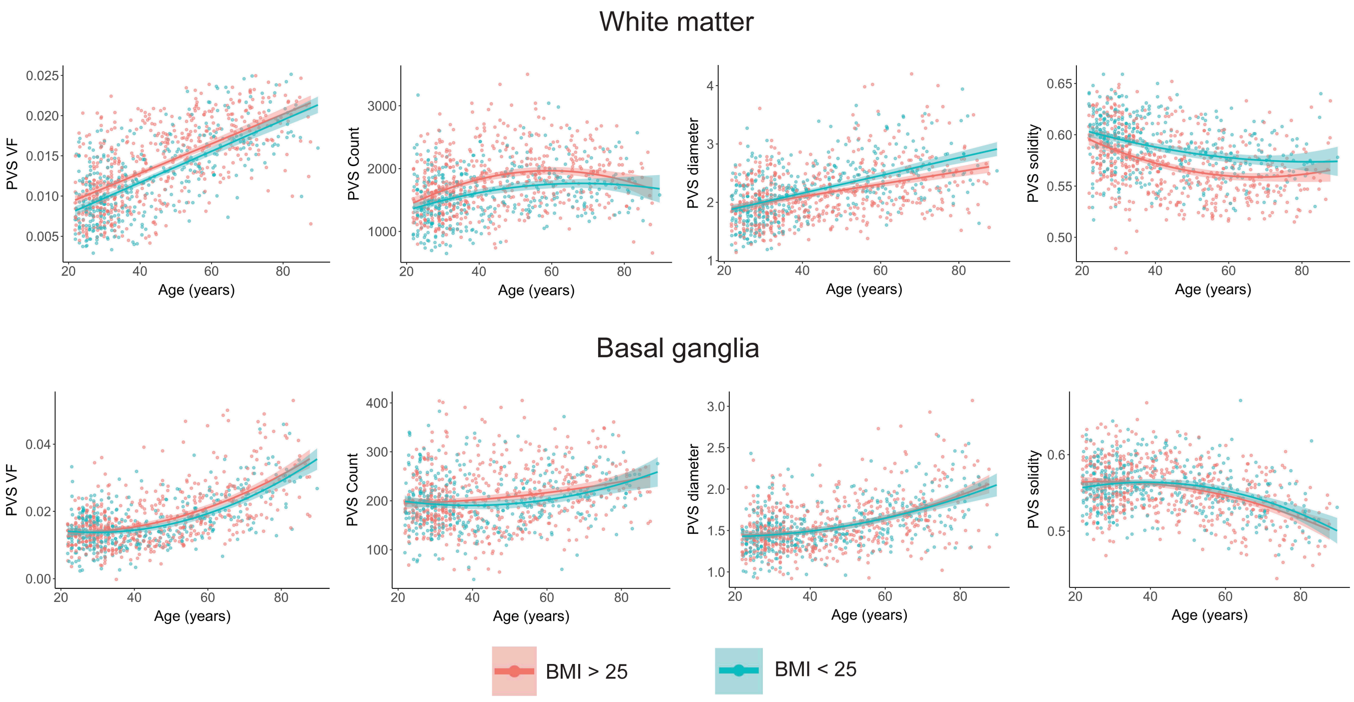


**Fig S4.** Correlations between BMI and PVS morphology in the white matter (top) and basal ganglia (bottom). Subjects over the age of 21 years of age were stratified into high BMI (>25) and low BMI (<25) cohorts. Data points reflect individual subjects and the best fit line with the estimated standard error are shown.
