## Supplementary material for "Brain perivascular space imaging across the human lifespan": SI Appendix, Figure S5

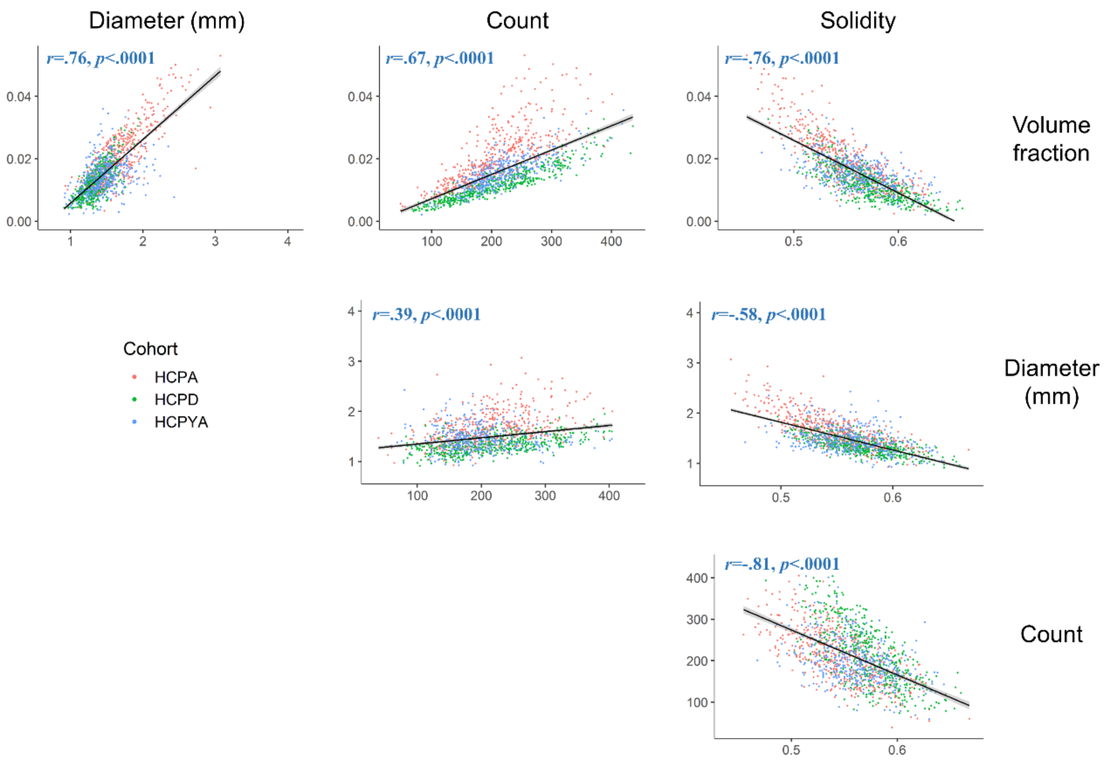


**Fig S4.** Morphological PVS correlations within BG. Data points are colored according to lifespan cohort. PVS morphological features within the BG were significantly correlated with one another.
