## Supplementary material for "Brain perivascular space imaging across the human lifespan": SI Appendix, Figure S6

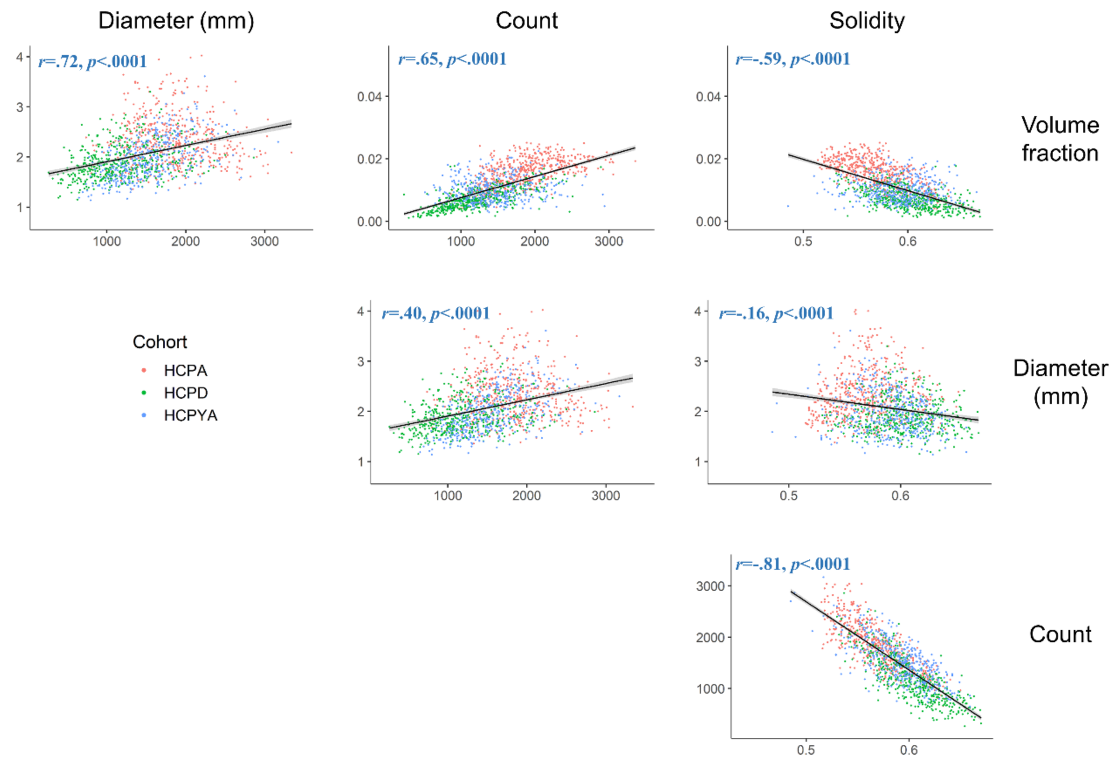


**Fig. S5.** Morphological PVS correlations within white matter. Data points are colored according to lifespan cohort. PVS morphological features within the white matter were significantly correlated with one another.
